# A Reproducible Pre-Transplant γδ T-cell Effector Signature Reveals Immune-Metabolic Programs Associated with Kidney Allograft Rejection

**DOI:** 10.64898/2026.10.31.763223

**Authors:** Seon Min Kim, Swastika Sur, Kelvin Koser, Silvia Pineda, Oriol Bestard, Troy Koyama, Thomas Hoffmann, Tara K. Sigdel, Marina Sirota, Joel Babdor, Matthew H. Spitzer, Minnie M. Sarwal

## Abstract

Despite optimized immunosuppression, acute kidney transplant rejection affects 10-20% of recipients within the first year, and no reliable pre-transplant molecular test exists to identify patients at high risk, independent of the donor HLA matching. Across three independent cohorts of kidney transplant recipients, profiled by bulk RNA-sequencing with T-cell receptor repertoire analysis, flow cytometry, CyTOF with paired metabolomics, we identified a reproducible depletion of γδ T cells associated with rejection: an elevated αβ/γδ T cell ratio in two post-transplant, biopsy-matched cohorts, and depletion of specific γδ T effector clusters in a third, longitudinally sampled cohort. In this third cohort, γδ T cell depletion was already present at the pre-transplant baseline in patients who subsequently developed rejection and remained detectable across the transplant course, indicating that it reflects recipient baseline immune biology rather than a consequence of graft injury. Longitudinal metabolomic profiling in this cohort identified time-dependent metabolic changes, including in amino acid metabolism that co-occur with the γδ signature and converge with it upon multi-omics integration. At the pre-transplant baseline, γδ T-cell marker expression alone showed exploratory discriminatory value for rejection risk (AUC = 0.69), improving marginally when integrated with metabolomic features (AUC = 0.71); larger prospective studies will be needed before clinical translation.

## 1 Introduction

Kidney transplantation remains the preferred treatment for end-stage renal disease, providing superior survival and quality of life compared with long-term dialysis(1). Despite advances in immunosuppressive therapy, allograft rejection remains a major cause of graft dysfunction and premature graft loss(2). Current surveillance strategies rely heavily on serum creatinine, donor-specific antibodies, and allograft biopsy, yet these approaches incompletely capture other determinants of rejection risk, including non-HLA antigens and pre-transplant immune activation signatures(3,4). Consequently, there remains substantial interest in identifying immune states associated with rejection risk before overt graft injury develops and in defining biomarkers that directly reflect the underlying biology of transplant outcomes. To date, however, most biomarker development in kidney transplantation has centered on post-transplant monitoring of biofluids, where transcriptomic, proteomic, and metabolomic profiling of blood and urine has advanced our understanding of injury-associated immune changes after engraftment(5). Pre-transplant immune states remain comparatively unexplored. Our group recently has shown that pediatric renal allograft recipients who go on to develop rejection already have a distinct pre-transplant B cell repertoire, with greater diversity and increased clonal expansion at baseline, and expanded clones that persist after rejection. Also, other molecular and functional cellular assays tracking donor-reactive T cells have also shown the relevance of preformed alloimmune memory driving transplant rejection(6–8). These pre-transplant findings were proposed as candidate markers for stratifying rejection risk before transplant, independent of donor source, HLA mismatch, and immunosuppression, although their predictive performance at the individual patient level remains modest(9).

Among the immune populations implicated in transplantation, γδ T cells represent a biologically distinct and incompletely understood lymphocyte lineage. Unlike conventional αβ T-cells, γδ T cells recognize stress-associated ligands, phosphoantigens, and other non-classical targets that are independent of MHC restriction, enabling rapid responses to injury, infection, and inflammation. Despite representing only a minor fraction of circulating T cells, γδ T cells are enriched in peripheral tissues and play important roles in immune surveillance, tissue homeostasis, and repair(10–12). Human γδ T cells are phenotypically and functionally heterogeneous, encompassing subsets with cytotoxic, pro-inflammatory, regulatory, and tissue protective properties that vary in their developmental origin and tissue context(13,14). This has complicated efforts to define a unifying role for γδ T cells in transplantation. In transplantation, γδ T cells have been linked to protective anti-viral responses(15,16), alloimmune injury and graft protection, with evidence for pathogenic roles in kidney allograft rejection and tolerance-associated signatures in liver transplantation(17–20). However, whether pre-transplant γδ T-cell states are associated with future kidney transplant outcomes has not been established. Addressing this question may provide insight into recipient immune predisposition before exposure to the allograft and immunosuppression.

To date, most studies of γδ T cells in kidney transplantation have focused on post-transplant observations, targeted subset analysis, or graft tissue findings. High-dimensional characterization of peripheral γδ T cells states before transplantation and their relationship to subsequent clinical outcomes has remained unexplored. Immune function is constrained by nutrient availability, and this link is particularly relevant for γδ T cells, whose recognition of phosphoantigens reflects sensing of metabolic intermediates rather than peptide antigens(21). γδ T cell subsets also carry distinct intrinsic metabolic programs linked to their effector functions(22), and nutrient availability more broadly shapes T cell proliferation, effector function, and lineage fate(23–26). Whether these metabolically-linked programs in γδ T cells are altered before transplantation, or predispose to eventual rejection, is unknown.

Resolving these states requires measurement at commensurate resolution. Mass cytometry quantifies dozens of phenotypic and functional markers per cell, enabling characterization of immune populations that population-level assays and routine clinical monitoring cannot distinguish(27,28). Metabolomics provides a functional readout of systemic biological functioning that biopsy and creatinine, respectively invasive and imprecise, do not capture(29). Applied together, the two modalities provide a comprehensive, holistic view of the systemic immune functioning, in which a metabolite signature becomes interpretable when anchored to the immune phenotype driving it.

Here we examined the relationship between γδ T cell states and transplant outcome across three independent cohorts of adult kidney allograft recipients. We hypothesized that distinct γδ T cell populations would be associated with rejection susceptibility and that joint immune and metabolic profiling would situate these populations within the broader biology underlying allograft outcomes. We find that the γδ T cells compartment is depleted in recipients who subsequently develop rejection, independent of donor, reproducibly across all three cohorts, and we characterized its relationship to systemic metabolic programs linked to rejection risk.

## 2 Materials and Methods

The complete experimental design spanning Cohorts 1-3 is illustrated in Figure 1.

**Figure 1.**
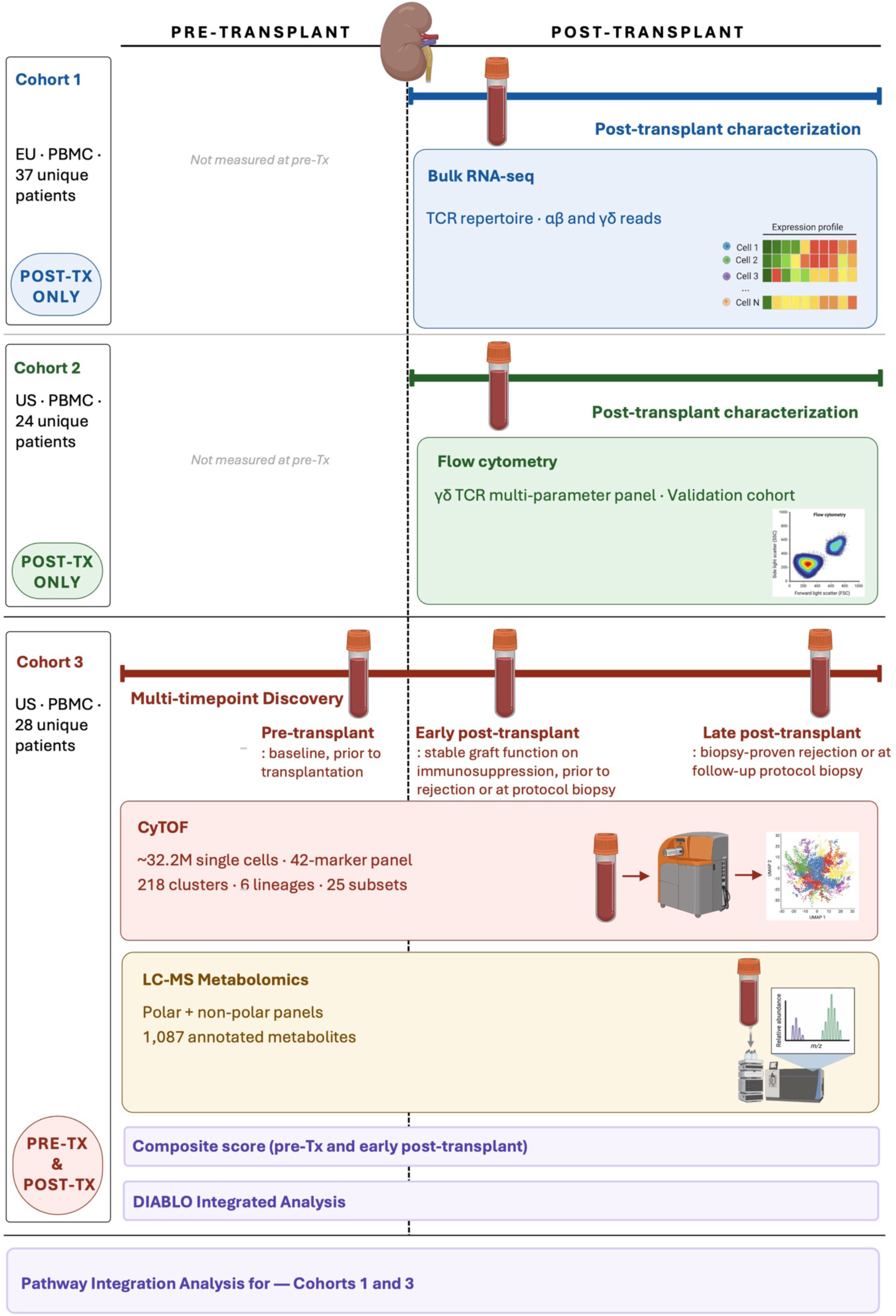
Study design. Three cohorts of kidney transplant recipients were profiled across two continents. Cohort 1 (EU, 37 patients) and Cohort 2 (US, 24 patients) provided post-transplant PBMCs for post-transplant characterization, analyzed by bulk RNA-seq and flow cytometry respectively; Cohort 2 served as an independent validation cohort. Cohort 3 (US, 28 patients) provided PBMCs collected at multiple timepoints before and after transplantation for multi-timepoint discovery, profiled by CyTOF (42-marker panel, ∼32.2M single cells, 218 clusters) and LC-MS plasma metabolomics (1,087 metabolites, polar and non-polar panels), with subsequent multi-omics integration (DIABLO) and pathway analysis across Cohorts 1 and 3. Dots indicate sample collection timepoints; kidney icon denotes transplant event (Day 0). For Cohort 3, samples were collected at three timepoints: pre-transplant (baseline, prior to transplantation and no immunosuppression), early post-transplant (stable graft function on immunosuppression, prior to rejection in acute rejection patient group or at 6 month protocol biopsy in stable patients), and late post-transplant (at biopsy-proven rejection in acute rejection patients or at follow-up protocol biopsy in NR patients). Cohort 3 outcomes (AR, NR) correspond to the AMR/TCMR/STA classification used in Cohort 1 and 2, with AR encompassing both AMR and TCMR and NR corresponding to STA. AMR, antibody-mediated rejection; AR, acute rejection; NR, non-rejection; PBMC, peripheral blood mononuclear cell; STA, stable; TCMR, T cell-mediated rejection.

### 2.1 Ethics statement

The studies involving humans were approved by the institutional review boards of Bellvitge University Hospital (Barcelona, Spain), the University of California, San Francisco, and the participating centers of the NIAID Clinical Trials in Organ Transplantation consortium. The studies were conducted in accordance with the local legislation and institutional requirements. The participants provided their written informed consent to participate in this study.

### 2.2 Cohort 1: RNA-SEQ

Bulk RNA-seq was performed on peripheral blood collected at the time of allograft biopsy from Cohort 1 patients with antibody-mediated rejection (AMR, n=12), T cell-mediated rejection (TCMR, n=13), or stable allografts (STA, n=12), as previously described(30).

#### 2.2.1 Data Processing and Quality Control

First, we used the TOIL publicly available pipeline to analyze the total number of reads, base quality score, and possible contamination of reads from adapters(31). All 37 samples passed the first two of these tests as they all possessed greater than 20 million total reads (the accepted minimum) and a quality score greater than 30 (the acceptable minimum; 40 is the maximum). Although the pipeline did detect overrepresentation of certain sequences in several samples, the fact that no adapter failures were detected seems to indicate that this abnormality is due to the sequence in question being highly biologically relevant (and thus overexpressed) rather than an error in the data collection.

#### 2.2.2 Immune Repertoire Analysis

For the immune repertoire analysis of the RNA-seq, we solely utilized the MiXCR pipeline(32).

The data was aligned to the Homo Sapiens reference genome, allowing for the partial alignment parameter; the inclusion of partial alignments ensures that incomplete reads are not discarded which allows for their reconstruction into complete reads later on in the process.

We then began round two of partial alignments (reconstructing incomplete reads from incomplete ones) followed by a single round of alignment extension, a step which reassembled incomplete CDR3 regions into completed CDR3 regions.

The last step before exporting the data was to assemble the clonotype data. Two immune cells are considered clones of each other if and only if their CDR3 sequences are of identical length and of 90% similarity. If these conditions are satisfied among two aligned reads, they are grouped together into a single clonotype, a group of immune cells likely representing products of clonal selection (a process by which immune cells coming into contact with antigens proliferate in order to develop the body’s immune response).

Finally, the data was exported into R where we conducted an ANOVA test on basic immune repertoire features (total reads, total clones, and reads/clones of IGH, IGK, IGL, TRA, TRB, TRD, and TRG) in addition to the α, β/γ, δ ratio(33). In addition to the data being sufficiently homoscedastic to justify the usage of an ANOVA test, we applied a non-parametric test (Kruskal-Wallis test) to ensure that the lack of normality was not an artifact.

#### 2.2.3 Gene Expression Analysis & Integration

The next stage of the RNA-seq data analysis began with the generation of a counts matrix representing the sample’s gene expression. To obtain the counts matrix, we used the STAR tool to align the filtered RNA-seq data to a comprehensive gene annotation of the CHR region (http://www.gencodegenes.org/releases/23.html)(34). This process allowed us to separate the immune repertoire genes (unmapped genes) from the coding genes used for the differential gene expression analysis (mapped genes). We then followed steps similar to other RNA-seq studies such as Liu et al. in order to fill in the count matrix with the gene expression information(35). This counts matrix was exported to the R environment where we utilized the DESeq2 package to normalize the matrix with the rlog transformation using the appropriate function(36). The last quality control step taken was to filter the counts matrix for an adjusted p-value < 0.05 and a | log 2 fold change | > 1. In order to ensure the normality of the data, we produced a PCA plot of the counts matrix (**Figure 2B**). With the samples being fairly uniformly spread out, we detected no observable batch effect or confounding factors.

**Figure 2.**
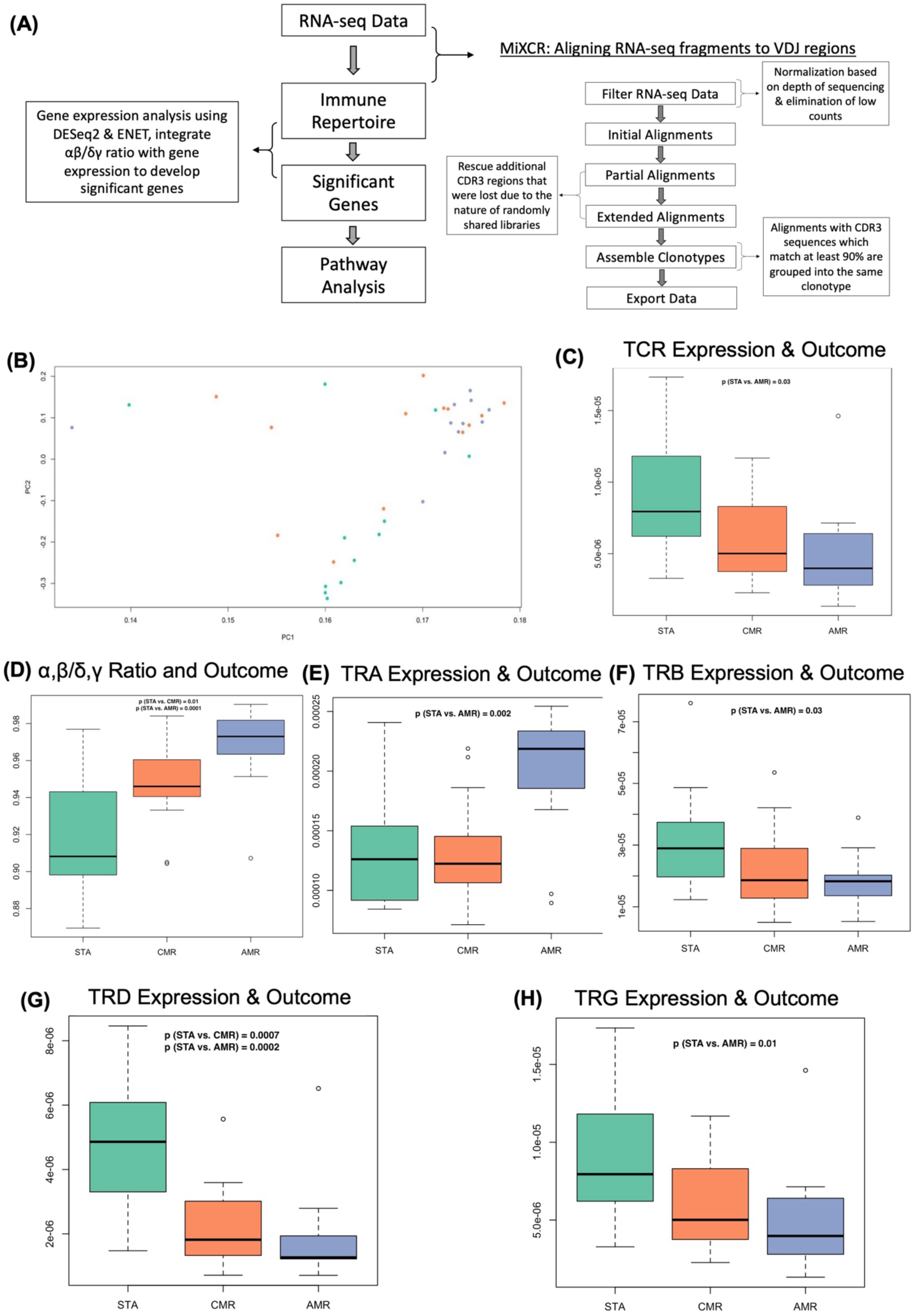
Overall workflow of RNA-seq analysis (Cohort 1), including the analytical pipeline and evaluation of total and subset-specific T-cell receptor (TCR) expression. In healthy individuals, the TCR repertoire is composed predominantly of αβ T cells (∼95%) with a smaller γδ T-cell population (∼5%), providing the baseline for comparison of relative αβ and γδ TCR expression across samples. **(A)** Detailed workflow from RNA-seq data processing, immune repertoire extraction, gene expression and pathway analysis including the steps in the MiXCR computational pipeline. **(B)** PCA plot of the normalized and quality-controlled gene expression counts matrix. **(C)** Boxplot results of TCR expression by transplant outcome. TCR expression is defined as number of reads mapping to TCR divided by total reads to normalize the expression to account for depth of sequencing. **(D)** Boxplot results for the classification by transplant outcome using α, β/γ, δ ratio. Vertical axis is percentage of all TCRs which are either TRA or TRB. **(E)** Boxplot results of α-TCR or TRA expression by transplant outcome. **(F)** Boxplot results of β-TCR or TRB expression by transplant outcome. (**G)** Boxplot results of δ -TCR or TRD expression by transplant outcome. (**H)** Boxplot results of γ -TCR or TRG expression by transplant outcome. Statistical significance was determined by one-way ANOVA after confirming homoscedasticity, with the Kruskal–Wallis test used to confirm robustness to non-normality. Pairwise comparisons were performed using t-tests. AMR, antibody-mediated rejection; TCMR, T cell-mediated rejection; STA, stable. Significance was determined by one-way ANOVA (data met homoscedasticity assumptions), with Kruskal-Wallis testing confirming robustness to non-normality.

We then applied a penalized regression method, Elastic NET (ENET), to find genes associated with the immune repertoire in the samples. We used the alpha tuning examples presented in Zou, H. and Hastie, T(37). The model attempted to select from the pool of all significantly differentially expressed genes (genes remaining after the filtering steps described above) those which were significantly associated with the α, β/γ, δ ratio.

#### 2.2.4 Pathway Analysis

Lastly, we introduced the significant genes from the ENET model into the **EnrichR and Reactome** tools in order to provide a biological interpretation(38–40). We did this separately for the downregulated and upregulated genes from the ENET model.

### 2.3 Cohort 2: Flow Cytometry

#### 2.3.1 Flow Cytometry

PBMCs were isolated by Ficoll method, and frozen in fetal bovine serum (FBS) with 10% Dimethyl sulfoxide (DMSO) for future use. On the day of the experiment, quick thaw followed by 5 min centrifugation at 350g and discard the supernatant. Resuspend the pellet in 100ul of cell staining buffer (BioLegend; Cat. No. 420201) then add 5ul of Human Fc Receptor Blocking Solution (TruStain FcX™, BioLegend Cat. No. 422301) and incubate it at room temperature for 20min. Human TruStain contains specialized human IgG that bind to Fc receptors, and is useful for reducing non-specific immunofluorescent staining. Next, we added appropriately conjugated fluorescent primary antibodies at predetermined optimum concentrations, and incubated in dark on ice for 30 min. The primary antibodies that we used are anti-human CD3 Antibody (PE/Cyanine7, Cat#317334), CD4 Antibody (Alexa Fluor® 700, Cat# 317426), CD8 Antibody (APC, Cat#344722) at a concentration of 1:1000, and TCR γ/δ Antibody (Alexa Fluor® 488, Cat#107512) at a concentration of 1:50. After the incubation period, add twice the volume of cell staining buffer (BioLegend; Cat. No. 420201) and centrifuge at 350g for 5min. Discard the supernatant. Repeat this step twice. Finally resuspend the cell pellet in 0.5ml of cell staining buffer (BioLegend; Cat. No. 420201). Just before sorting, add 5µl (0.25µg)/million cells of 7-AAD Viability Staining Solution (BioLegend, Cat. No. 420403), and incubate on ice for 5 min and perform fluorescence activated cell sorting (FACS). We included negative control and a Fluorescence minus One (FMO) control per run. FMO is a tube of cells stained with all fluorochromes used in the experiment except one. A multi-color immunofluorescent experiment has one FMO control for each fluorochrome. It helps to determine the cut-off point between background fluorescence and positive populations in multi-color immunofluorescent experiments. The sample size for each group in our experiment are stable (n=10), TCMR (n=8), and AMR (n=6), paired with the allograft biopsy. We classified the cells into 2 categories - αβ and γδ. Cells which are CD3-positive, γδ-negative and double or single positive for CD4 and CD8 are in αβ category. Whereas cells that are CD3-positive, γδ-positive and double or single negative for CD4 and CD8 are in γδ category.

### 2.4 Cohort 3: Immune-metabolomic Profiling

#### 2.4.1 Study Cohort

A total of 28 kidney transplant recipients were enrolled in a longitudinal observational study. Patients were stratified into two clinical groups according to biopsy-confirmed rejection status: acute rejection (n = 9) and non-rejection (n = 19).

Longitudinal peripheral blood samples were collected at three clinically defined timepoints: pre-transplantation, early post-transplantation, and late post-transplantation. The early post-transplantation timepoint was collected prior to biopsy confirmation of rejection status, while the late post-transplantation timepoint corresponded to the period following biopsy-proven rejection in acute-rejection patients. In stable patients, the post-transplantation timepoints were defined by a 6-month protocol biopsy, at which no rejection was identified. Peripheral blood mononuclear cells (PBMCs) were isolated from each collected sample and stored for subsequent analysis. A total of 80 PBMC samples were available across all patients and timepoints and were carried forward into the analyses described below.

#### 2.4.2 CyTOF Sample Collection and Processing

PBMCs were isolated from peripheral blood and cryopreserved according to a standardized protocol prior to batch processing. Cryopreserved samples were subsequently thawed and prepared for single-cell mass cytometry (CyTOF) analysis using an established workflow.

Pooled, barcoded samples were stained with a 42-marker antibody panel targeting markers of immune cell lineage identity and functional state, and data were acquired on a CyTOF mass cytometry platform.

##### 2.4.2.1 Mass-tag antibody conjugation and antibody cocktail preparation for mass cytometry

Mass cytometry antibodies were prepared using the MaxPAR antibody conjugation kit (Standard Bio Tools) according to the manufacturer’s instructions. Following conjugation, antibodies were diluted at concentrations ranging from 0.2 and 6 mg/mL in Candor PBS Antibody Stabilization solution (Candor Bioscience GmbH, Wangen, Germany) supplemented with 0.02% NaN3 and stored at 4°C. Each antibody was titrated to identify optimal staining concentrations using human PBMCs. A surface staining antibody cocktail and an intracellular antibody cocktail made of all the 2 types of target were prepared for the staining of all batches of the study.

##### 2.4.2.2 Mass cytometry sample preparation

Briefly, PBMCs were incubated immediately after isolation with a viability staining solution containing 50uM of cisplatin (Sigma-Aldrich) for 1 min at room temperature and fixed in a solution containing 1.6% Paraformaldehyde (PFA) (Electron Microscopy Science) and cryopreserved in a solution containing 0.5% BSA and 10% dimethyl sulfoxide (DMSO). Fixed cells were later thawed and incubated with benzonase nuclease (Sigma-Aldrich), permeabilized with a solution containing 0.2% saponin, barcoded by incubation with a combination of custom palladium metal stable isotopes for 15 minutes at room temperature, and pooled in batches of 20 samples. Fc receptors were blocked using Human TruStain FcX (Biolegend) and cells were stained with custom metal ion-labeled antibody cocktails. Extracellular staining was performed at room temperature for 30 minutes on a shaker set on 90 RPM. Intracellular staining was then performed using eBioscience Permeabilization Buffer (Invitrogen) at 4°C for 60 minutes on a shaker set on 90 RPM. Cells were later incubated with Cell-ID Intercalator-Ir (Standard Biotools) at 0.0625 uM and 4% PFA overnight at 4°C. Before running, cells were resuspended in cell acquisition solution (Standard Biotools) that contained EQ Four Element Calibration Beads (Standard Biotools). Except for permeabilization steps and final steps, washes were performed using a cell staining solution containing PBS, BSA and EDTA or PBS.

##### 2.4.2.3 Mass cytometry data acquisition

Mass cytometry sample pools were diluted in Cell Acquisition Solution containing bead standards to approximately 1 x 10^ 6 cells/mL and then analyzed on a Helios mass cytometer (Standard Biotools) equilibrated with Cell Acquisition Solution. A minimum of 10 x 10^ 6 cell events were collected for each barcoded set of samples at an event rate of 400 events/second.

#### 2.4.3 CyTOF Data Analysis

Bead standard data normalization and de-barcoding of pooled samples into their respective conditions was performed using the premessa R package, with removal of normalization beads, dead cells, and doublets. Live, single-cell events were arcsinh-transformed using a cofactor of 5 and batch-corrected using cyCombine (v0.2.19). Unsupervised clustering and UMAP-based dimensionality reduction were performed using cyclone, which implements FlowSOM (v2.0.0; 15×15 grid) for clustering and the uwot R package (v0.1.10) for UMAP embedding, yielding 225 single-cell clusters across 80 PBMC samples. After post-annotation quality filtering to remove artifact and doublet clusters, the remaining 218 clusters were manually annotated using canonical lineage markers into 25 immune subsets across 6 lineages. All 42 markers from the panel were used as input for clustering and dimensionality reduction.

#### 2.4.4 CyTOF Statistical Analysis

To identify immune subsets distinguishing patient groups (acute rejection vs. stable graft), cell-subset frequencies were treated as compositional data and transformed using the centered log-ratio (CLR) transformation (with a pseudocount of 1e-5 to handle zeros) using the compositions R package. Partial Least Squares Discriminant Analysis (PLS-DA) was then applied to CLR-transformed frequencies using the mixOmics R package, and features were ranked by Variable Importance in Projection (VIP) scores. Top-ranked features were validated using Wilcoxon rank-sum tests with Benjamini-Hochberg FDR correction (FDR-adjusted p < 0.05). This analysis was performed independently at each of the three timepoints.

#### 2.4.5 LC-MS Metabolomics

##### 2.4.5.1 Sample extraction and acquisition

Frozen plasma samples (200 µl) were thawed on ice and extracted with 800 µl of 1:1 acetonitrile:methanol containing 2-amino-3-bromo-5-methylbenzoic acid (1 mg/ml) and a mixture of ¹³C,¹⁵N-labeled amino acids (Sigma Aldrich, cat. #767964; 100 pg/µl) as internal standards. Samples were vortexed, held at 4 °C for 10 min, and centrifuged at 14,000 rpm for 30 min at 4 °C to pellet precipitated protein. A 250 µl aliquot of supernatant was transferred to vials and analyzed immediately.

Untargeted high-resolution mass spectrometry was performed on a Sciex ExionLC UPLC coupled to a Sciex TripleTOF 6600+ mass spectrometer. A 5 µl aliquot was injected onto a Kinetex F5 column (2.6 µm, 100 Å, 2.1 × 150 mm; Phenomenex, cat. #00F-4723-AN). Mobile phase A was water with 0.1% formic acid and mobile phase B was acetonitrile with 0.1% formic acid, at a constant flow rate of 200 µl/min. The 20-minute gradient was: 0.2% B (0 min), 0.2% B (2 min), 98% B (14 min), 98% B (16 min), 0.2% B (16.1 min), 0.2% B (20 min). Data were acquired separately in positive (ISVF = 5500 V) and negative (ISVF = −4500 V) ionization modes using data-dependent acquisition, across m/z 50–1000, with the top 20 most abundant ions selected for MS/MS and a total cycle time of 0.650 s. Collision energy was 35 V with a spread of 15 V; DP = 40, EP = 10, CXP = 15, GS1 = 60, GS2 = 80, CUR = 35, IRD = 66, IRW = 24, TEM = 400 °C. Peak detection and processing were performed using SCIEX OS (v2.0.0.45330) and MarkerView (v1.4).

##### 2.4.5.2 Data Processing

Features were annotated against the KEGG compound database; unannotated features were discarded. Features with >50% missingness across samples were removed, data were log2-transformed, and remaining missing values were imputed as half the minimum observed value per feature. Batch effects were corrected using limma::removeBatchEffect(). After preprocessing, 628 annotated polar features and 459 annotated non-polar features were retained for downstream analysis.

##### 2.4.5.3 Metabolomics Analysis

To identify discriminatory metabolites between patient groups, Random Forest classifiers (500 trees) were trained using nested cross-validation: leave-one-out cross-validation (LOOCV) was used as the outer loop, with VSURF feature selection performed within each training fold to prevent information leakage. Classifier performance was quantified using AUC computed from out-of-fold predictions. Pathway-level enrichment was performed by mapping metabolites to KEGG compound IDs and running GSEA-based enrichment using the fgsea R package, with metabolites ranked by log2 fold change between groups, producing normalized enrichment scores (NES) and BH-adjusted p-values. All analyses were performed independently at three timepoints (pre-transplant, early post-transplant, late post-transplant).

#### 2.4.6 Multi-omics Integration

Two complementary integrative analyses were performed on cohort 3 samples profiled by both CyTOF and untargeted LC-MS.

##### 2.4.6.1 Composite biomarker panel

To build a biomarker panel discriminating acute rejection from stable allograft at the pre-transplant timepoint, the per-patient median expression of selected markers (CD161, CD45RA, gdTCR, CD3, CD27) within γδ T cells was combined with metabolites from the polar and non-polar LC-MS panels. The composite panel was evaluated using nested leave-one-out cross-validation (LOOCV) with Random Forest classifiers (500 trees), in which VSURF feature selection was performed on the polar and non-polar metabolomics data within each training fold to prevent information leakage. Performance was compared against a baseline model using only γδ T cell marker expression. AUC was computed from out-of-fold predictions, with 95% confidence intervals derived from 2,000 bootstrap iterations.

##### 2.4.6.2 DIABLO multi-block integration

To identify maximally correlated features across CyTOF and metabolomics platforms, DIABLO (block sparse PLS-DA) from the mixOmics R package was applied independently at each of the three timepoints (pre-transplant, early post-transplant, late post-transplant). Prior to integration, metabolomics features were restricted to biologically annotated compounds; near-zero-variancefeatures were removed, duplicate annotations were collapsed to a single representative feature, and four adduct-derived duplicate annotations identified during manual QC were excluded, yielding 536 metabolite features per timepoint. Two latent components were extracted from annotated CyTOF clusters and metabolomics features, with a design matrix off-diagonal value of 0.1. Feature selection retained 10 CyTOF clusters and 20 metabolites on component 1, and 5 CyTOF clusters and 10 metabolites on component 2. Model performance was estimated using 3-fold cross-validation with 5 repeats (mixOmics perf()); balanced error rate (BER) computed using centroid distances is reported for each block and component. Feature selection was repeated within each training fold.

## 3 Results

To investigate peripheral blood biomarkers of acute kidney allograft rejection, we assembled three independent cohorts spanning two continents, allowing for inclusion of patients with varying demographics and ethnicity (**Figure 1**). Demographics and clinical characteristics for each cohort are provided in **Tables 1-3**. Cohort 1 (Bellvitge University Hospital, Barcelona University, IDIBELL, Barcelona, Spain, EU, n=37) and Cohort 2 (UCSF, USA, n=24) were cross-sectional and sampled post-transplant only, profiled by bulk RNA-seq with TCR repertoire analysis and flow cytometry, respectively. Cohort 3 (US, n=28) included paired pre-transplant and post-transplant samples profiled by CyTOF and LC-MS metabolomics, with sampling at pre-transplant, early post-transplant, and late post-transplant timepoints.

**Table 1.**
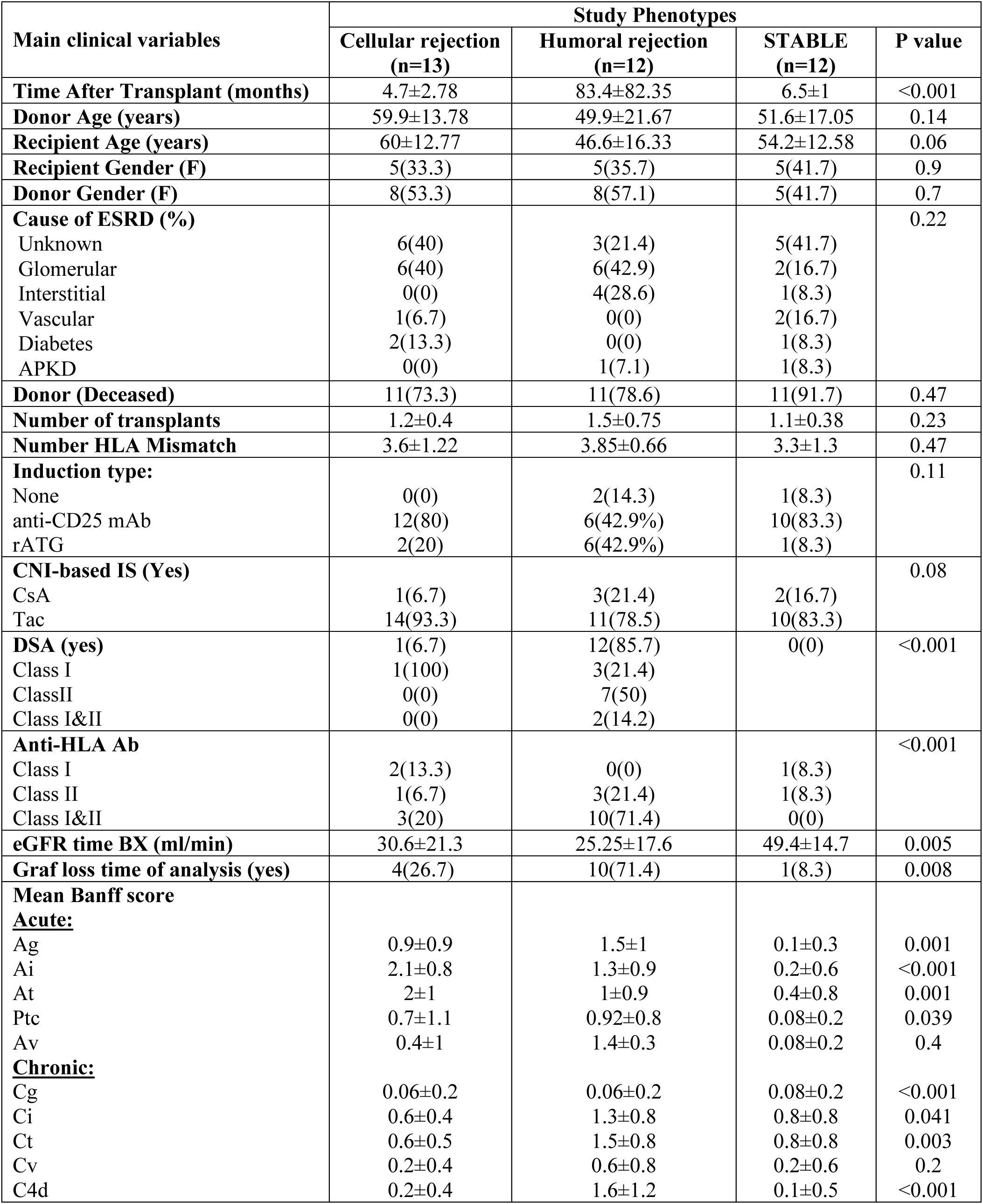
Clinical and Demographic Characteristics of Cohort 1.

### 3.1 Depletion of γδ TCR chain transcripts associates with rejection

Bulk RNA-seq was performed in cohort 1 on PBMCs isolated from peripheral blood of biopsy-matched antibody-mediated rejection (AMR, n=12), T cell-mediated rejection (TCMR, n=13), and stable (STA, n=12) samples **(Table 1)**. Immune repertoire analysis of post-transplant PBMC RNA-seq yielded TCR and BCR clonotypes across all seven loci (**Supplementary Figure 1, Figure 2A**). Principal component analysis showed that the samples did not show outcome-based clustering or systematic separation by technical variables, confirming that no dominant confounding factor drove global transcriptomic variance in this cohort (**Figure 2B**).

Overall TCR expression was significantly higher in STA compared to AMR (p=0.03; **Figure 2C**), indicative of a global reduction in T cell receptor transcript abundance in rejection. The αβ/δγ TCR ratio was the most discriminatory immune repertoire feature across outcomes: both TCMR and AMR showed a significantly elevated ratio compared to STA (TCMR: p=0.01; AMR: p=0.0001; **Figure 2D**), reflecting a relative depletion of γδ T cells in rejection. Given its strong discriminatory performance across both rejection subtypes, the αβ/γδ ratio was selected as the immune repertoire feature for downstream integration with gene expression analysis. Consistent with the elevated αβ/δγ ratio, individual αβ TCR chains showed opposing directionality to γδ chains across outcomes. T cell receptor alpha (TRA) expression was significantly higher in AMR than STA (p=0.002; **Figure 2E**), with no significant difference between STA and TCMR. T cell receptor beta (TRB) expression was significantly lower in AMR compared to STA (p=0.03; **Figure 2F**), with a non-significant trend toward reduction in TCMR. In humans, αβ T cells constitute approximately 95% and γδ T cells approximately 5% of peripheral T cells under homeostatic conditions. The elevated TRA in AMR alongside reduced TRB suggests a skewed αβ compartment rather than uniform T cell loss. The strongest and most consistent rejection signal was observed in the γδ TCR chains. Expression of TRD that encodes the δ chain of the γδ TCR was significantly lower in both TCMR (p=0.0007) and AMR (p=0.0002) compared to STA (**Figure 2G**), representing the most statistically significant individual chain difference in this cohort. Expression of TRG, encoding the g chain of the gd TCR, was significantly lower in AMR compared to STA (p=0.01; **Figure 2H**), with a non-significant trend in TCMR. Together, the concordant reduction of TRD and TRG across both rejection subtypes, at both the read and clonotype level (**Supplementary Figure 1**), establishes γδ TCR chain depletion as the primary immune repertoire signature distinguishing stable allografts from rejection in Cohort 1.

Next, we wanted to visualize the gene expression levels of the significant coding genes in each sample by transplant outcome. The heatmap (**Supplementary Figure 2**) represents the significant differential gene expression between AMR and STA. There were 1716 significantly differentially expressed genes, out of which 1506 were upregulated and 210 were downregulated in the AMR samples. However, no significant differential expression of genes was observed between STA and TCMR.

### 3.2 Genes associated with the αβ/δγ ratio, enriched in allograft rejection and antigen presentation pathways

To identify the transcriptional programs associated with the αβ/γδ ratio, we regressed the ratio on the full coding gene expression matrix using an elastic net (ENET) model, selecting genes independently of transplant phenotype. The ENET model (α, β/γ, δ *ratio* ∼ *gene expression matrix)* applied to the entire list of significant coding genes found a total of 81 genes significantly associated with the ratio as shown in the heatmap (**Figure 3A**). Out of these 81 genes significantly associated with the ratio, 23 were downregulated, and 58 were upregulated. Although the samples did not cluster perfectly by phenotype, the signature of these genes associated with the α, β/γ, and δ ratio was informative. This was anticipated, as the model selected genes based on their association with the αβ/γδ ratio rather than with transplant outcome. 81 Genes from ENET Model are shown in **Supplemental Table 1**.

**Figure 3.**
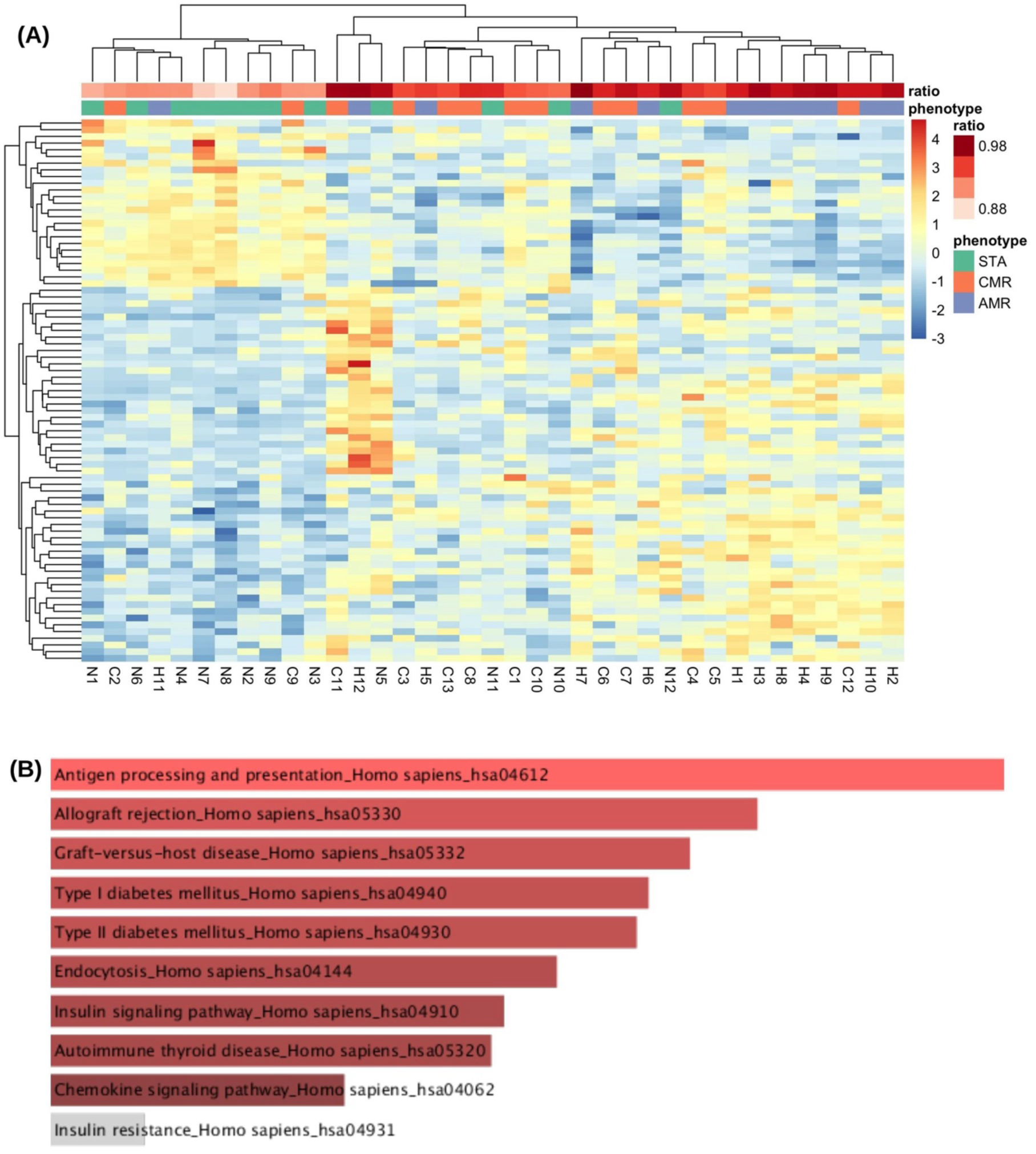
Dataset 1: genes associated with α,β/δ,γ ratio and their corresponding significantly enriched pathways. **(A)** Heatmap visualizing results of ENET model (α, β/γ, δ *ratio* ∼ *gene expression matrix*). The 81 genes (23 downregulated and 58 upregulated in rejection outcomes) are shown on the vertical axis while the horizontal access represents individual samples (color-coded by transplant outcome). The value of the α, β/γ, δ ratio is also depicted on the horizontal access. **(B)** Pathway Analysis by Enrichr identified the most significant pathways included allograft rejection (adjusted p-value = 0.001) together with antigen processing and presentation (unadjusted p-value = 1.25 10 * -5 ) and graft-vs-host disease (unadjusted p-value = 2.24 10 * -5). Gene ontology (STRING) enrichment analysis identified significant overrepresentation of pathways related to T-cell activation and programmed cell death. Reactome pathway analysis further highlighted enrichment of Interleukin-10 signaling and activation of complement components C3 and C5, suggesting coordinated regulation of adaptive immune responses, immune suppression, and complement-mediated inflammatory processes.

After carrying out pathway analysis by Enrichr, we found that the most significant pathways were allograft rejection (adjusted p-value = 0.001), antigen processing and presentation (unadjusted p-value = 1.25 × 10^-5^) and graft-vs-host disease (unadjusted p-value = 2.24 × 10^-5^) (**Figure 3B**).

### 3.3 γδ T cell depletion distinguishes rejection subtypes in an independent US cohort

In order to validate our prior findings, we applied a multiparameter flow cytometry panel to analyze cryopreserved PBMCs from an independent US cohort (Cohort 2; STA n=10, TCMR n=8, AMR n=6; **Table 2**) paired with biopsy-confirmed allograft outcomes. The gating strategy is shown in **Figure 4A**. In a representative stable patient (Specimen_001-7), γδ T cells comprised a mean of 0.22% ± 0.14% of double-negative T cells in the STA group, consistent with the known low peripheral frequency of this population. Quantification across outcome groups (**Figure 4B-D**) showed that γδ T cell frequency was significantly higher in TCMR than in both STA and AMR (p<0.05 for each). STA and AMR did not differ from one another. This indicates that γδ T cell expansion is specific to TCMR. αβ T cell frequency showed the converse pattern. It was significantly elevated in AMR relative to both STA and TCMR (p<0.001 for each), while STA and TCMR did not differ. This indicates that αβ T cell expansion is instead specific to AMR. Because each rejection type is marked by expansion of a different T cell compartment, the αβ/γδ ratio separated all three groups (STA vs TCMR, p<0.05; STA vs AMR, p<0.01; TCMR vs AMR, p<0.0001). TCMR was lowest, driven by the γδ increase, and AMR was highest, driven by the αβ increase. This shows that the ratio does not simply flag the presence of rejection. It flags which T cell compartment is expanding, and therefore which mode of rejection is occurring. Critically, γδ T cell frequency alone distinguished TCMR from AMR, a clinically important discrimination that standard immunological markers frequently fail to achieve. These findings independently validate in a geographically distinct cohort the γδ T cell depletion signal first identified by RNA-seq in Cohort 1 and extend it by demonstrating that the magnitude of γδ T cell loss differs between rejection subtypes, with AMR showing the most severe depletion.

**Figure 4.**
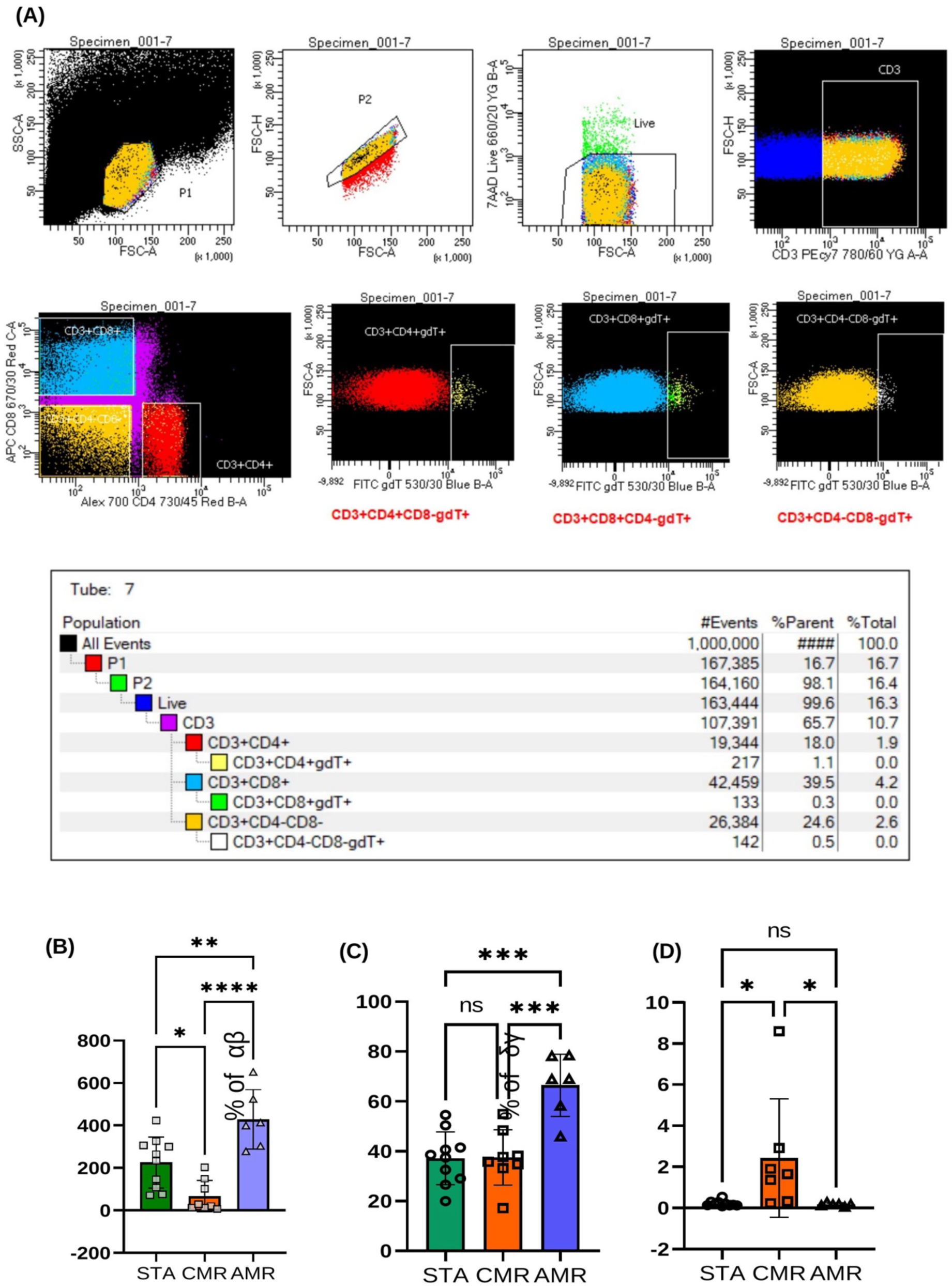
Dataset 2: Gating strategy and γδ T cell quantification in Cohort 2 (n=24). Quantification of TCR subset frequencies across transplant outcomes: stable (STA, n=10), T cell-mediated rejection (TCMR, n=8), and antibody-mediated rejection (AMR, n=6), paired with allograft biopsy. **(A)** Representative flow cytometry gating hierarchy from a stable allograft patient (Specimen_001-7). PBMCs were stained with CD3-PE/Cy7, CD4-Alex700, CD8-APC, and TCRγδ-FITC, with 7-AAD viability exclusion. Sequential gates were applied as follows: P1, lymphocyte scatter gate (FSC-A vs SSC-A); P2, singlet discrimination (FSC-H vs FSC-A); Live, 7-AAD-negative viable cells; CD3, T cell gate (FSC-H vs CD3-PE/Cy7). Within CD3+ T cells, subsets were defined by CD4 and CD8 co-expression followed by TCRγδ-FITC: CD3+CD4+gdT+ (red), CD3+CD8+CD4-gdT+ (cyan), and CD3+CD4-CD8-gdT+ (orange, canonical γδ T cells). Population statistics from a representative sample are shown in the accompanying hierarchy table (Tube 7). The canonical γδ T cell gate (CD3+CD4-CD8-gdT+) represented 0.5% of the CD3+CD4-CD8-double-negative fraction in this sample. The same gating hierarchy was applied uniformly to all Cohort 2 samples. Gate boundaries for TCRγδ-FITC were set using a fluorescence-minus-one (FMO) control run in parallel for each sample. **(B)** Bar plot results for the classification by transplant outcome using α, β/ δ, γ ratio at the protein level. **(C)** CD3-positive, γδ-negative and double or single positive for CD4 and CD8 are in αβ category. Bar plot shows % of αβ T-cells for the classification by transplant outcome. **(D)** Cells that are CD3-positive, γδ-positive and double or single negative for CD4 and CD8 are in γδ category. Bar plot shows % of γδ T-cells for the classification by transplant outcome. Results from One way ANOVA & Tukey’s multiple comparison test. *p < 0.05, **p <0.01, ***p <0.001, ****p <0.0001.

**Table 2.** Clinical and demographic characteristics of Cohort 2.

| **Main Clinical Variables** | **Cellular rejection<br>(n=10)** | **Humoral rejection<br>(n=6)** | **STABLE<br>(n=11)** | **P value** |
|---|---|---|---|---|
| **Time After Transplant (months)** | 41.5±55.7 | 94.4±66.0 | 7.4±2.4 | <0.001 |
| **Recipient Age (years)** | 44.1±14.4 | 43.2±6.9 | 41.3±13.0 | 0.81 |
| **Recipient Gender (F)** | 5(50.0) | 2(33.3) | 3(27.3) | 0.55 |
| **Cause of ESRD (%)**<br>Unknown<br>Glomerular<br>Interstitial<br>Vascular<br>Diabetes<br>APKD<br>Other | 0(0.0)<br>3(30.0)<br>2(20.0)<br>1(10.0)<br>2(20.0)<br>1(10.0)<br>1(10.0) | 0(0.0)<br>1(16.7)<br>0(0.0)<br>1(16.7)<br>0(0.0)<br>3(50.0)<br>1(16.7) | 1(9.1)<br>4(36.4)<br>1(9.1)<br>0(0.0)<br>3(27.3)<br>1(9.1)<br>1(9.1) | 0.54 |
| **Donor Type**<br>DD<br>LRD<br>LURD<br>DAMMI | 5(50.0)<br>2(20.0)<br>2(20.0)<br>1(10.0) | 2(33.3)<br>2(33.3)<br>2(33.3)<br>0(0.0) | 5(45.5)<br>1(9.1)<br>5(45.5)<br>0(0.0) | 0.65 |
| **Number of Transplants** | 1.1±0.3 | 1.0±0.0 | 1.1±0.3 | 0.74 |
| **Induction Type**<br>anti-CD25 mAb<br>rATG | 3(33.3)<br>6(66.7) | 2(66.7)<br>1(33.3) | 6(54.5)<br>5(45.5) | 0.50 |
| **CNI-based IS**<br>Tac<br>Other (Belatacept) | 8(80.0)<br>2(20.0) | 5(100.0)<br>0(0.0) | 11(100.0)<br>0(0.0) | 0.18 |
| **Clinical BPAR**<br>No<br>Borderline<br>Yes | 0(0.0)<br>0(0.0)<br>10(100.0) | 0(0.0)<br>2(33.3)<br>4(66.7) | 11(100.0)<br>0(0.0)<br>0(0.0) | <0.001 |
| **eGFR at Time of Bx (mL/min)** | 53.1±24.7 | 53.3±16.4 | 79.6±20.6 | 0.04 |
| **Mean Banff Score**<br>Acute:<br>Ag<br>Ai<br>At<br>Ptc<br>Av<br>Chronic:<br>Cg<br>Ci<br>Ct<br>Cv<br>C4d | 0.90±0.99<br>1.90±0.99<br>1.60±1.07<br>1.11±1.17<br>0.40±0.70<br>0.10±0.32<br>1.00±0.94<br>1.30±0.67<br>1.60±1.26<br>0.00±0.00 | 2.83±0.41<br>0.33±0.52<br>1.00±0.63<br>1.67±0.52<br>0.17±0.41<br>2.17±0.98<br>1.33±0.82<br>1.50±0.55<br>1.67±1.21<br>2.75±0.50 | 0.00±0.00<br>0.00±0.00<br>0.00±0.00<br>0.00±0.00<br>0.00±0.00<br>0.00±0.00<br>0.00±0.00<br>0.00±0.00<br>0.00±0.00<br>0.00±0.00 | <0.001<br><0.001<br><0.001<br><0.001<br>0.16<br><0.001<br>0.00<br><0.001<br>0.00<br><0.001 |

### 3.4 Pre-transplant γδ T-cell clusters associated with rejection phenotypes

To further evaluate the γδ T cell signal observed in earlier cohorts, we profiled Cohort 3, an independent multi-omics cohort on contemporaneous samples comprising 28 patients with acute rejection (AR; n=9) or non-rejection (NR; n=19; **Table 3**), with a 42-marker CyTOF panel. Using unsupervised clustering via FlowSOM, we identified 218 distinct immune clusters, spanning 6 lineages and 25 subsets. UMAP analysis allowed us to visualize the distribution of these populations in two-dimensional space (**Figure 5A**).

**Table 3.**
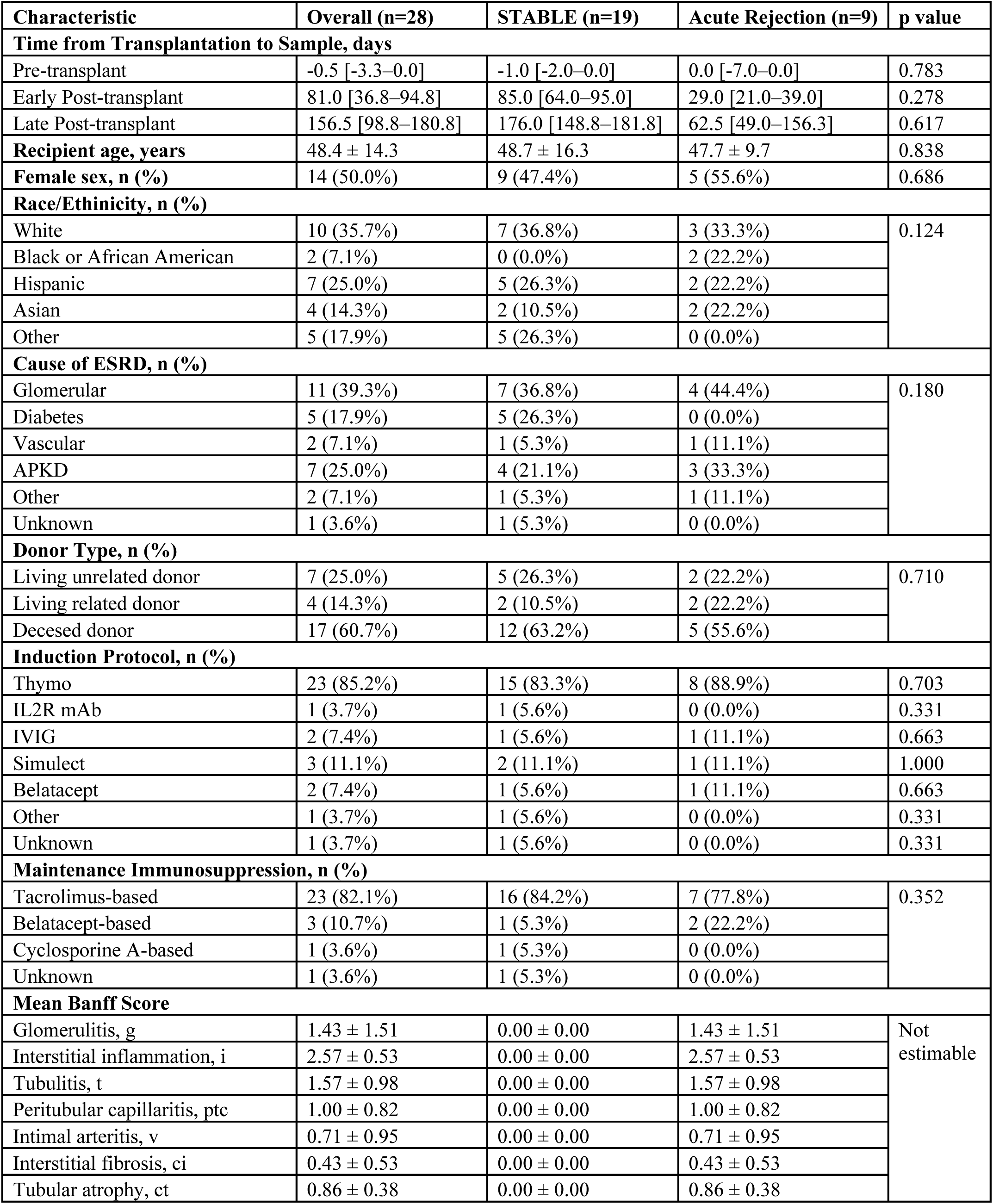
Clinical and demographic characteristics of Cohort 3.

| Characteristic | Overall (n=28) | STABLE (n=19) | Acute Rejection (n=9) | p value |
|---|---|---|---|---|
| **Time from Transplantation to Sample, days** |  |  |  |  |
| Pre-transplant | -0.5 [-3.3–0.0] | -1.0 [-2.0–0.0] | 0.0 [-7.0–0.0] | 0.783 |
| Early Post-transplant | 81.0 [36.8–94.8] | 85.0 [64.0–95.0] | 29.0 [21.0–39.0] | 0.278 |
| Late Post-transplant | 156.5 [98.8–180.8] | 176.0 [148.8–181.8] | 62.5 [49.0–156.3] | 0.617 |
| **Recipient age, years** | 48.4 ± 14.3 | 48.7 ± 16.3 | 47.7 ± 9.7 | 0.838 |
| **Female sex, n (%)** | 14 (50.0%) | 9 (47.4%) | 5 (55.6%) | 0.686 |
| **Race/Ethinicity, n (%)** |  |  |  |  |
| White | 10 (35.7%) | 7 (36.8%) | 3 (33.3%) | 0.124 |
| Black or African American | 2 (7.1%) | 0 (0.0%) | 2 (22.2%) |  |
| Hispanic | 7 (25.0%) | 5 (26.3%) | 2 (22.2%) |  |
| Asian | 4 (14.3%) | 2 (10.5%) | 2 (22.2%) |  |
| Other | 5 (17.9%) | 5 (26.3%) | 0 (0.0%) |  |
| **Cause of ESRD, n (%)** |  |  |  |  |
| Glomerular | 11 (39.3%) | 7 (36.8%) | 4 (44.4%) | 0.180 |
| Diabetes | 5 (17.9%) | 5 (26.3%) | 0 (0.0%) |  |
| Vascular | 2 (7.1%) | 1 (5.3%) | 1 (11.1%) |  |
| APKD | 7 (25.0%) | 4 (21.1%) | 3 (33.3%) |  |
| Other | 2 (7.1%) | 1 (5.3%) | 1 (11.1%) |  |
| Unknown | 1 (3.6%) | 1 (5.3%) | 0 (0.0%) |  |
| **Donor Type, n (%)** |  |  |  |  |
| Living unrelated donor | 7 (25.0%) | 5 (26.3%) | 2 (22.2%) | 0.710 |
| Living related donor | 4 (14.3%) | 2 (10.5%) | 2 (22.2%) |  |
| Decesed donor | 17 (60.7%) | 12 (63.2%) | 5 (55.6%) |  |
| **Induction Protocol, n (%)** |  |  |  |  |
| Thymo | 23 (85.2%) | 15 (83.3%) | 8 (88.9%) | 0.703 |
| IL2R mAb | 1 (3.7%) | 1 (5.6%) | 0 (0.0%) | 0.331 |
| IVIG | 2 (7.4%) | 1 (5.6%) | 1 (11.1%) | 0.663 |
| Simulect | 3 (11.1%) | 2 (11.1%) | 1 (11.1%) | 1.000 |
| Belatacept | 2 (7.4%) | 1 (5.6%) | 1 (11.1%) | 0.663 |
| Other | 1 (3.7%) | 1 (5.6%) | 0 (0.0%) | 0.331 |
| Unknown | 1 (3.7%) | 1 (5.6%) | 0 (0.0%) | 0.331 |
| **Maintenance Immunosuppression, n (%)** |  |  |  |  |
| Tacrolimus-based | 23 (82.1%) | 16 (84.2%) | 7 (77.8%) | 0.352 |
| Belatacept-based | 3 (10.7%) | 1 (5.3%) | 2 (22.2%) |  |
| Cyclosporine A-based | 1 (3.6%) | 1 (5.3%) | 0 (0.0%) |  |
| Unknown | 1 (3.6%) | 1 (5.3%) | 0 (0.0%) |  |
| **Mean Banff Score** |  |  |  |  |
| Glomerulitis, g | 1.43 ± 1.51 | 0.00 ± 0.00 | 1.43 ± 1.51 | Not estimable |
| Interstitial inflammation, i | 2.57 ± 0.53 | 0.00 ± 0.00 | 2.57 ± 0.53 |  |
| Tubulitis, t | 1.57 ± 0.98 | 0.00 ± 0.00 | 1.57 ± 0.98 |  |
| Peritubular capillaritis, ptc | 1.00 ± 0.82 | 0.00 ± 0.00 | 1.00 ± 0.82 |  |
| Intimal arteritis, v | 0.71 ± 0.95 | 0.00 ± 0.00 | 0.71 ± 0.95 |  |
| Interstitial fibrosis, ci | 0.43 ± 0.53 | 0.00 ± 0.00 | 0.43 ± 0.53 |  |
| Tubular atrophy, ct | 0.86 ± 0.38 | 0.00 ± 0.00 | 0.86 ± 0.38 |  |

**Figure 5.**
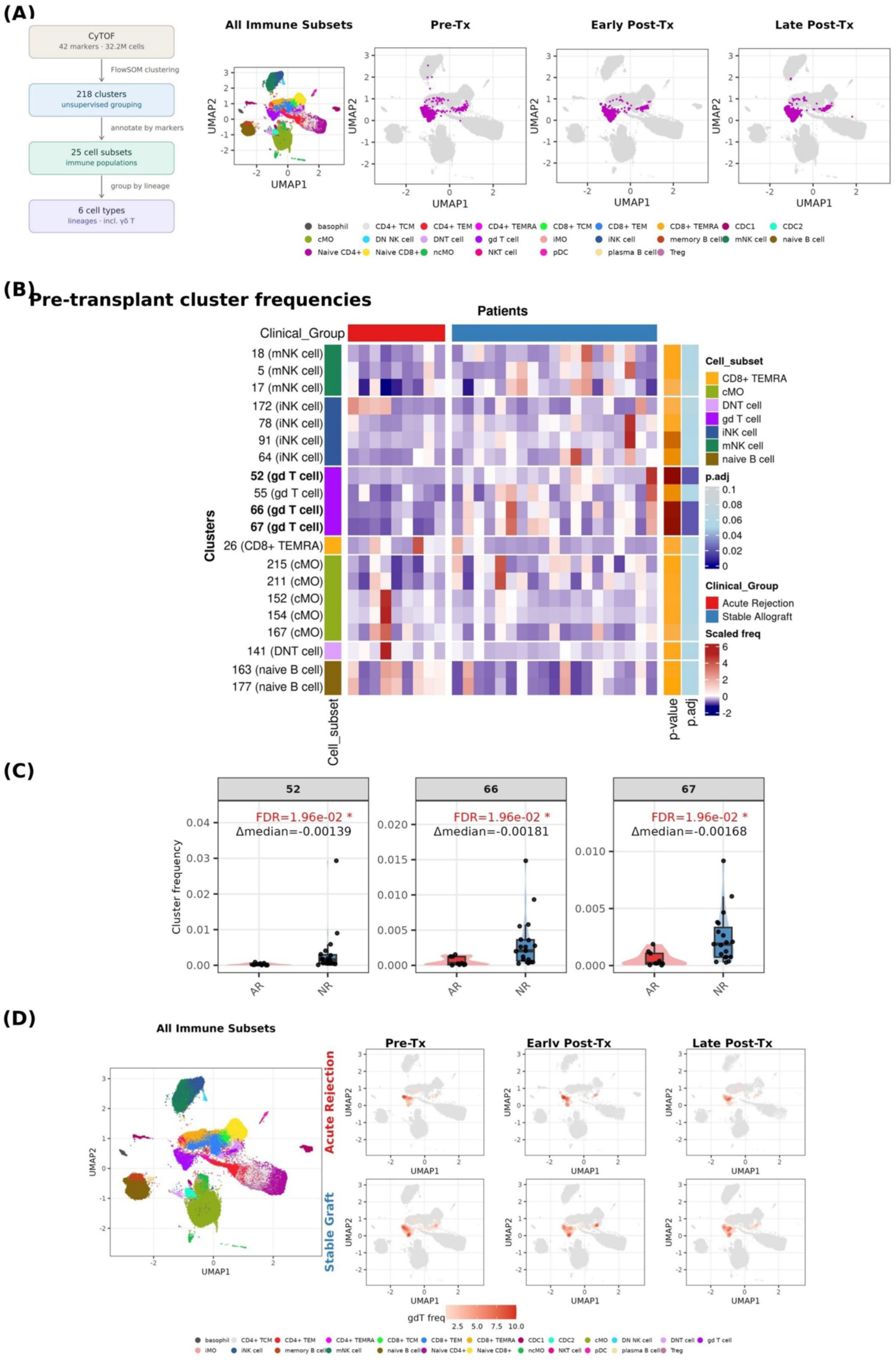
High-dimensional mass cytometry reveals pre-transplant depletion of effector γδ T cells in patients who develop acute rejection. **(A)** UMAP visualization of 32.2 million single cells from Cohort 3 (n=28 patients), with 218 FlowSOM clusters resolved into 25 immune subsets. Cells are shown across three timepoints — pre-transplant, early post-transplant, and late post-transplant — to capture the evolving immune landscape surrounding transplantation. **(B)** Heatmap of z-score normalized cluster frequencies in pre-transplant samples, stratified by clinical outcome (acute rejection, red; non-rejection, blue). Clusters were ranked by PLS-DA VIP scores and the top 20 selected features are shown. γδ T cell clusters 52, 66, and 67 (purple sidebar) are consistently depleted in acute rejection patients. **(C)** Pre-transplant frequencies of γδ T cell clusters 52, 66, and 67 are significantly lower in patients who subsequently developed acute rejection compared to non-rejectors (FDR=1.96×10⁻², Wilcoxon rank-sum test with BH correction applied within the 20 PLS-DA–prioritized clusters). **(D)** γδ T cell frequencies projected onto the UMAP across timepoints, separately for patients who developed acute rejection and those with stable grafts. γδ T cells are notably enriched in stable graft patients prior to transplantation.

We first examined whether predefined immune subsets at the pre-transplant baseline could discriminate clinical outcomes. Pre-transplant immune architecture appeared broadly similar between groups at the global level, suggesting any outcome-associated differences were subtle in nature rather than reflecting overt compositional reorganization (**Supplementary Figure 3A**). Pre-transplant UMAP feature plots illustrated qualitative differences in the spatial distribution of several immune subsets between outcome groups, with γδ T cells, naive B cells, and mature NK (mNK) cells appearing more abundant in non-rejection patients, and CD8+ Terminally differentiated Effector Memory re-expressing (TEMRA) and CD4+ central memory T cell (TCM) more prominent in certain acute rejection individuals (**Supplementary Figure 3C**). To identify which subsets most strongly separated outcome groups, immune subsets were ranked using Partial Least Squares Discriminant Analysis (PLS-DA), and the top-ranked subsets were carried forward for formal testing. Formal testing of the top PLS-DA-prioritized immune cell subsets showed no significant pre-transplant differences after multiple testing corrections (**Supplementary Figure 4A**). At early post-transplant, γδ T cells were significantly reduced in acute rejection patients after FDR correction, while other prioritized subsets showed nominal or non-significant differences (**Supplementary Figure 4B**).

We therefore asked whether higher-resolution phenotypic analysis might uncover signals obscured by population-level aggregation. FlowSOM clusters were ranked by PLS-DA variable importance in projection (VIP) scores, and the top 20 clusters were examined further. Because this ranking was informed by outcome labels, the resulting p-values are not valid for formal inference and are reported as exploratory. Within this set, differences concentrated in γδ T cell-annotated clusters, while no αβ T cell, NK, monocyte, or B cell cluster reached significance at this threshold (**Figure 5B, Supplementary Table 2**).

Three γδ T cell clusters (52, 66, and 67) were consistently depleted in patients who subsequently developed acute rejection relative to those with stable grafts (**Figure 5C**). Uniform Manifold Approximation and Projection (UMAP) overlays across all timepoints (pre-transplant, early post-transplant, late post-transplant) and outcome groups showed that these clusters localized to a compact, overlapping γδ T cell region within the immune landscape (**Figure 5D**), reflecting phenotypically similar γδ T cell states. Consistent with this, UMAP density overlays showed reduced occupancy of this region in acute rejection patients compared to non-rejectors **(Supplementary Figure 3B)**.

Cluster 55 was the fourth γδ T cell cluster among the top 20 but did not differ significantly by outcome. To characterize what distinguished clusters 52, 66, and 67 from cluster 55, we compared median expression across all 42 markers (**Supplementary Figure 5**). While all four clusters expressed canonical γδ T cell markers — CD3, CD45, and γδTCR — clusters 52, 66, and 67 showed relatively elevated expression of CD161 (the largest positive marker difference, at approximately 2.54 in arcsinh-transformed median expression), alongside higher expression of CD57, CD56, T-bet, and Ki67, broadly consistent with a cytotoxic, effector-memory-like phenotype. Conversely, the rejection-associated clusters showed markedly lower CD27 and CD45RA relative to cluster 55, suggesting a more terminally differentiated state with loss of a naive-associated surface phenotype. These patterns support annotation of clusters 52, 66, and 67 as antigen experienced effector γδ T cells, distinct from cluster 55. These comparisons remain descriptive given the modest sample size, but the phenotypic contrast is internally consistent across markers.

We examined the two most divergent markers, CD161 and CD45RA, in greater detail, contrasting the three significant clusters (52, 66, 67) with the non-significant cluster 55 at all three timepoints. Both markers differed significantly after BH correction at every timepoint (FDR < 0.05; **Supplementary Figure 5B**), with CD161 markedly enriched and CD45RA reduced in the significant clusters. This indicates that the phenotypic differences between significant and non-significant γδ clusters are reproducible across timepoints and are reflected at the marker-expression level. We note, however, that this comparison characterizes clusters that were defined by their differential abundance and does not by itself establish CD161 or CD45RA as population-level discriminators of rejection.

Collectively, these analyses indicate that outcome-associated signal in the γδ compartment resides in specific effector subpopulations rather than across all γδ T cells. Three γδ clusters depleted before transplantation in patients who subsequently developed acute rejection shared a cytotoxic, terminally differentiated phenotype marked by high CD161, CD57, CD56, and T-bet and low CD27 and CD45RA, distinguishing them from the non-associated γδ cluster 55. Pre-transplant depletion of these effector γδ subpopulations represents a candidate feature distinguishing patients who develop acute rejection from those with stable graft function, though replication in larger independent cohorts will be necessary to assess its robustness.

### 3.5 Timepoint-specific candidates distinguishing acute rejection

Cohort 3 was profiled with paired LC-MS plasma metabolomics alongside CyTOF, sampled at pre-transplant, early post-transplant, and late post-transplant timepoints, yielding 1,087 annotated metabolites across polar and non-polar panels. As circulating metabolites reflect systemic immune and metabolic state and are readily measurable in blood, we next examined whether metabolomic profiles could similarly distinguish acute rejection from stable graft outcomes across the transplant course. To do this, we applied a nested leave-one-out cross-validation (LOOCV) framework in which Variable Section Using Random Forests (VSURF)-based feature selection(41) was performed independently on the training partition at each fold, ensuring that the held-out sample contributed no information to feature selection. A Random Forest classifier was then trained on the VSURF-selected features and evaluated against the held-out sample, with this procedure repeated across all folds. This nested design was applied separately to polar and non-polar metabolomic datasets in Cohort 3.

To explore which metabolites were most consistently selected across folds, we applied a stability threshold of ≥50% of folds; all observations below this threshold should be regarded as hypothesis-generating given the small cohort size (**Figure 6**). At pre-transplant, D-pantethine and γ-butyrolactone were the stably selected features in the polar panel, each selected in ≥50% of folds and tending toward nominally higher levels in the rejection group (p = 0.016 and 0.041; FDR = 0.83). In the non-polar panel, 3-methylamino-L-alanine and δ-hexanolactone met the stability threshold at pre-transplant. At early post-transplant, 5-methyltetrahydrofolic acid (polar panel) and p-cresol (non-polar panel) were the only features meeting the ≥50% stability threshold, with both selected in nearly all cross-validation folds. At late post-transplant, loratadine met the stability threshold in the non-polar pane and likely reflects medication exposure rather than endogenous biology. Because feature stability varied across timepoints, these findings should be considered exploratory and require validation in independent cohorts. Overall, relatively few metabolites exceeded the predefined stability threshold, and the identities of stable figures differed across timepoints, suggesting that outcome associated metabolic signals were temporarily dynamic rather than driven by a common set of metabolites throughout the transplant time-course sampled.

**Figure 6.**
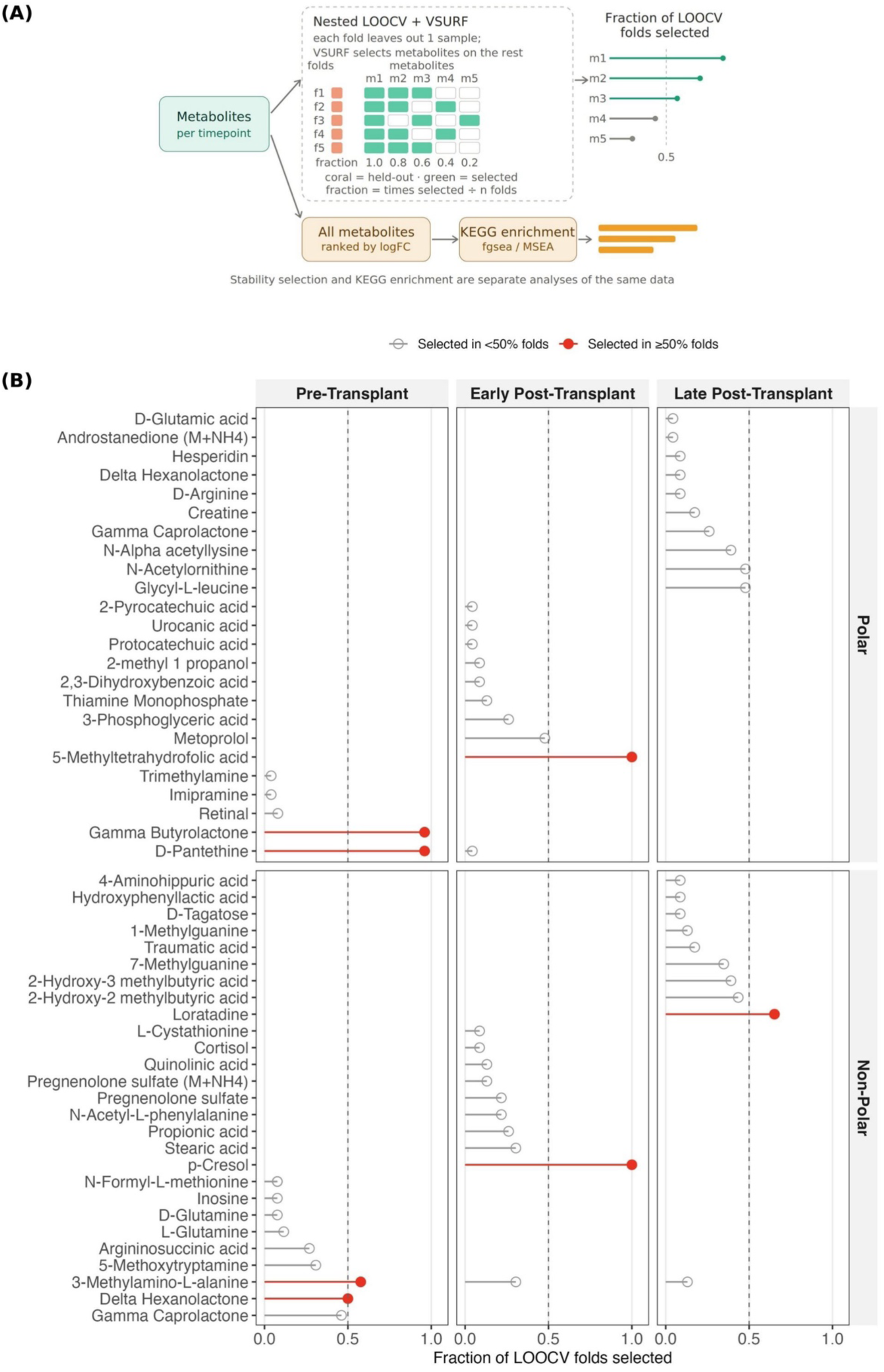
Stability of metabolite feature selection across cross-validation folds. **(A)** Schematic of the nested LOOCV and VSURF workflow used to calculate metabolite selection stability. In each fold, VSURF was performed on the training samples only, and selection frequency was defined as the fraction of LOOCV folds in which each metabolite was selected. **(B)** Fraction of LOOCV folds selected for metabolites across polar and non-polar datasets at each timepoint. Top metabolites are ranked by selection frequency. Metabolites selected in ≥50% of folds are shown in red; the dashed line marks the 50% threshold.

### 3.6 Timepoint-specific metabolic differences for acute rejection

To explore biological pathways associated with acute rejection, we performed KEGG(42) metabolite set enrichment analysis (MSEA) using fgsea(43) on the full ranked metabolite list at each timepoint, independent of the VSURF-based feature selection pipeline. Metabolites were ranked by log-fold change between groups and pathway enrichment was tested across KEGG pathway sets, with results filtered at FDR < 0.10 (**Figure 7**).

**Figure 7.**
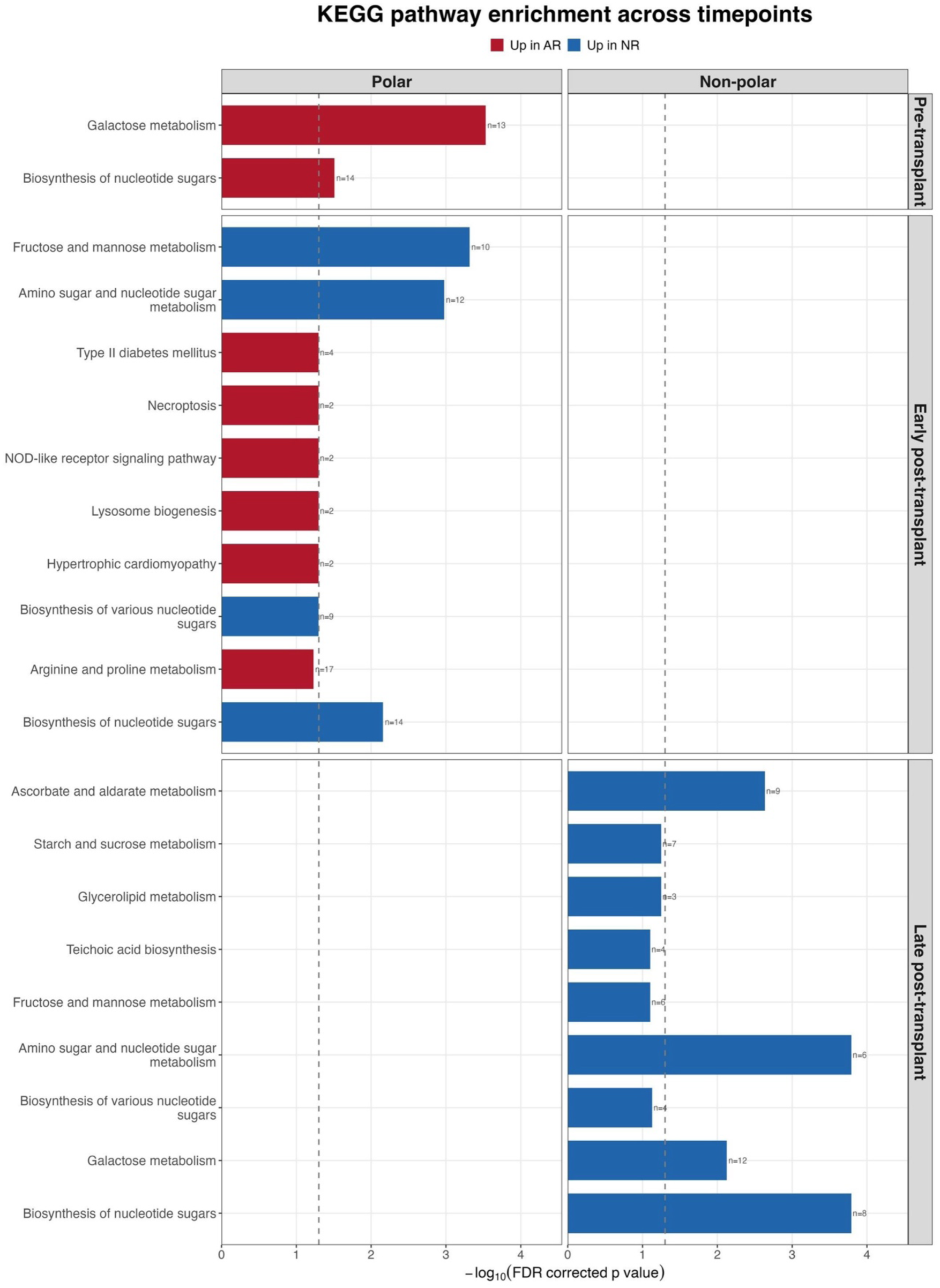
Longitudinal MSEA reveals AR-specific metabolic reprogramming. Metabolite Set Enrichment Analysis (MSEA) of KEGG pathways across polar and non-polar LC-MS panels at pre-transplant, early post-transplant, and late post-transplant timepoints. Bars represent −log₁₀(FDR-corrected p-value); dashed line indicates FDR = 0.05. n indicates the number of detected metabolites in each pathway set. Red bars denote pathways enriched in acute rejection (AR; NES > 0); blue bars denote pathways enriched in non-rejection (NR; NES < 0). No significant pathways were detected in the non-polar panel at late post-transplant.

At pre-transplant, two polar pathways were enriched in acute rejection patients at FDR < 0.10: galactose metabolism (n=13 metabolites) and biosynthesis of nucleotide sugars (n=14). No pathways met the FDR < 0.10 threshold in the non-polar fraction at this timepoint.

At early post-transplant, the polar fraction showed enrichment across multiple pathways. In acute rejection patients, these included arginine and proline metabolisms (n=17), biosynthesis of various nucleotide sugars (n=9), type II diabetes mellitus (n=4), necroptosis (n=2), NOD-like receptor signaling pathway (n=2), lysosome biogenesis (n=2), and hypertrophic cardiomyopathy (n=2). Because enrichment results based on very few mapped metabolites are unstable, we restricted interpretation to pathways represented by at least five metabolites. The pathways with n=2 above are therefore reported for completeness only. In stable graft patients, enriched polar pathways included fructose and mannose metabolism (n=10), amino sugar and nucleotide sugar metabolism (n=12), and biosynthesis of nucleotide sugars (n=14). No pathways met the FDR < 0.10 threshold in the non-polar fraction at this timepoint.

At late post-transplant, no pathways met the FDR < 0.10 threshold in the polar fraction. In the non-polar fraction, all enriched pathways were enriched in stable graft patients, including amino sugar and nucleotide sugar metabolism (n=6), biosynthesis of nucleotide sugars (n=8), galactose metabolism (n=12), ascorbate and aldarate metabolism (n=9), starch and sucrose metabolism (n=7), fructose and mannose metabolism, teichoic acid biosynthesis, biosynthesis of various nucleotide sugars (n=4), and glycerolipid metabolism (n=3).

Across timepoints, the most immunologically interpretable signal was enrichment of arginine and proline metabolism in acute rejection patients at the early post-transplant. Although the magnitude of this enrichment was modest, it was supported by a substantially larger metabolite set than other pathways meeting threshold at this timepoint (n=17 versus n=2–4), and arginine-related features were independently selected at the same timepoint in the integrative analysis described below. Arginine availability regulates T cell cell-cycle progression and effector function, and myeloid arginase activity depletes extracellular arginine with consequent suppression of proliferation and cytokine production(23,24), providing a plausible link between the metabolic and cellular findings in this cohort.

The remaining enriched pathways were dominated by carbohydrate and nucleotide sugar metabolism. Given that plasma carbohydrate profiles are sensitive to pre-analytical variation, and that the direction of enrichment for these pathways was not consistent across timepoints, being enriched in acute rejection at pre-transplant but in stable graft patients at late post-transplant, these are not interpreted further.

### 3.7 Combining γδ T cell and metabolomic features suggest complementary information for rejection risk assessment

Having identified outcome associated γδ T cell features and time-point specific metabolite candidates, we next asked whether combining these two modalities could improve risk stratification at the pre-transplant and at the early post-transplant timepoint. We constructed a composite classifier incorporating γδ T cell marker expression features alongside timepoint-specific VSURF-selected metabolites and evaluated its performance under nested leave-one-out cross-validation, with feature selection performed independently on the training partition at each fold to prevent data leakage.

γδ T cell marker expression at pre-transplant alone yielded modest discriminatory performance (AUC = 0.692, 95% CI: 0.496–0.887). VSURF-selected metabolites alone performed comparably (AUC = 0.647, 95% CI: 0.406–0.865), and the combined model produced a produced a marginal improvement (AUC = 0.714, 95% CI: 0.489–0.902; **Figure 8A**). At early post-transplant, metabolomic features alone yielded the highest point estimate (AUC = 0.733, 95% CI: 0.525–0.933), with the combined model performing similarly (AUC = 0.713, 95% CI: 0.467–0.900) and γδ T cell markers alone performing least well (AUC = 0.625, 95% CI: 0.375–0.842; **Figure 8B**). Given the substantial overlap in confidence intervals, these data do not establish that any model outperforms the others, and all should be considered exploratory given the sample size.

**Figure 8.**
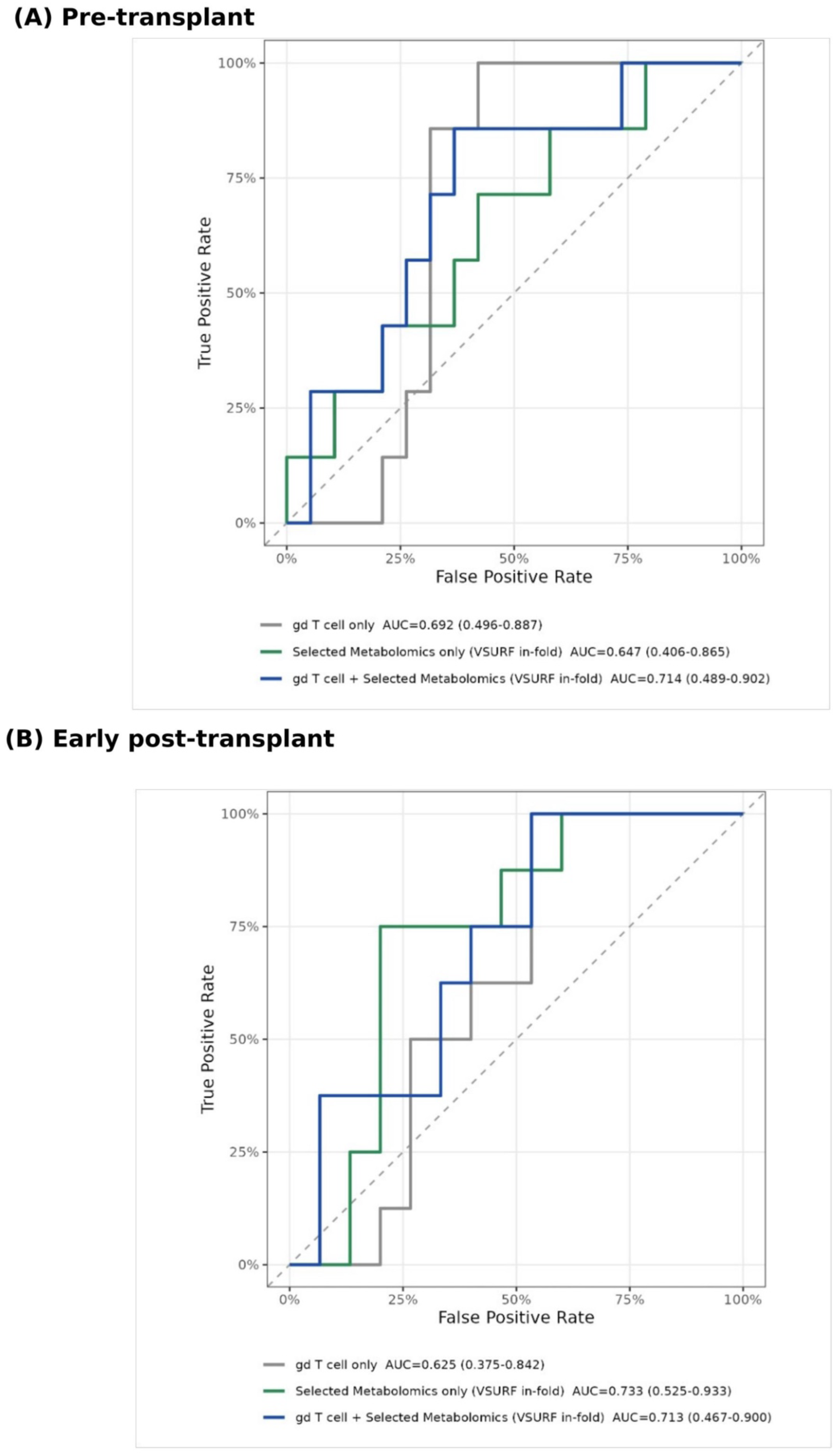
Predictive performance of γδ T cell and metabolomic features for acute rejection. ROC curves from nested LOOCV Random Forest models predicting acute rejection versus non-rejection at **(A) pre-transplant** and **B) early post-transplant** timepoints. Three models are compared: a γδ T cell-only baseline (grey), a model using VSURF-selected metabolomic features alone (green), and a combined model incorporating both γδ T cell and VSURF-selected metabolomic features (blue). Feature selection was performed within each fold using training data only to prevent data leakage.

### 3.8 DIABLO reveals time-dependent immune-metabolite associations across the transplant course

The preceding analyses examined the CyTOF and metabolomic data separately, which cannot reveal whether the immune and metabolic signals are related to one another. To ask whether γδ T cell features and metabolites co-vary in a coordinated manner, we applied DIABLO(44), a supervised multi-omics integration framework that identifies correlated features across data types while discriminating outcome groups. Features selected by DIABLO may therefore differ from those identified in the preceding individual analyses; such discordance is expected by design and reflects the distinct questions each analytical framework is optimized to answer.

To explore coordinated immune–metabolite patterns associated with acute rejection, multi-block integration of DIABLO was applied independently at each timepoint. Sample projections onto the first two latent components showed partial separation between acute rejection and stable graft patients in both blocks at all three timepoints (**Figure 9B**). Cross-validated classification performance was modest overall and varied by timepoint and block: balanced error rates were lowest at late post-transplant (metabolomics 0.255, CyTOF 0.345) and highest at early post-transplant, where CyTOF performance on the second component did not exceed chance (0.504). The metabolomics block outperformed the CyTOF block at every timepoint (**Supplementary Table 3**). In the supervised DIABLO latent space, the metabolomics block showed tighter group clustering than CyTOF across timepoints, suggesting that metabolite features contributed strongly to class separation this model.

**Figure 9.**
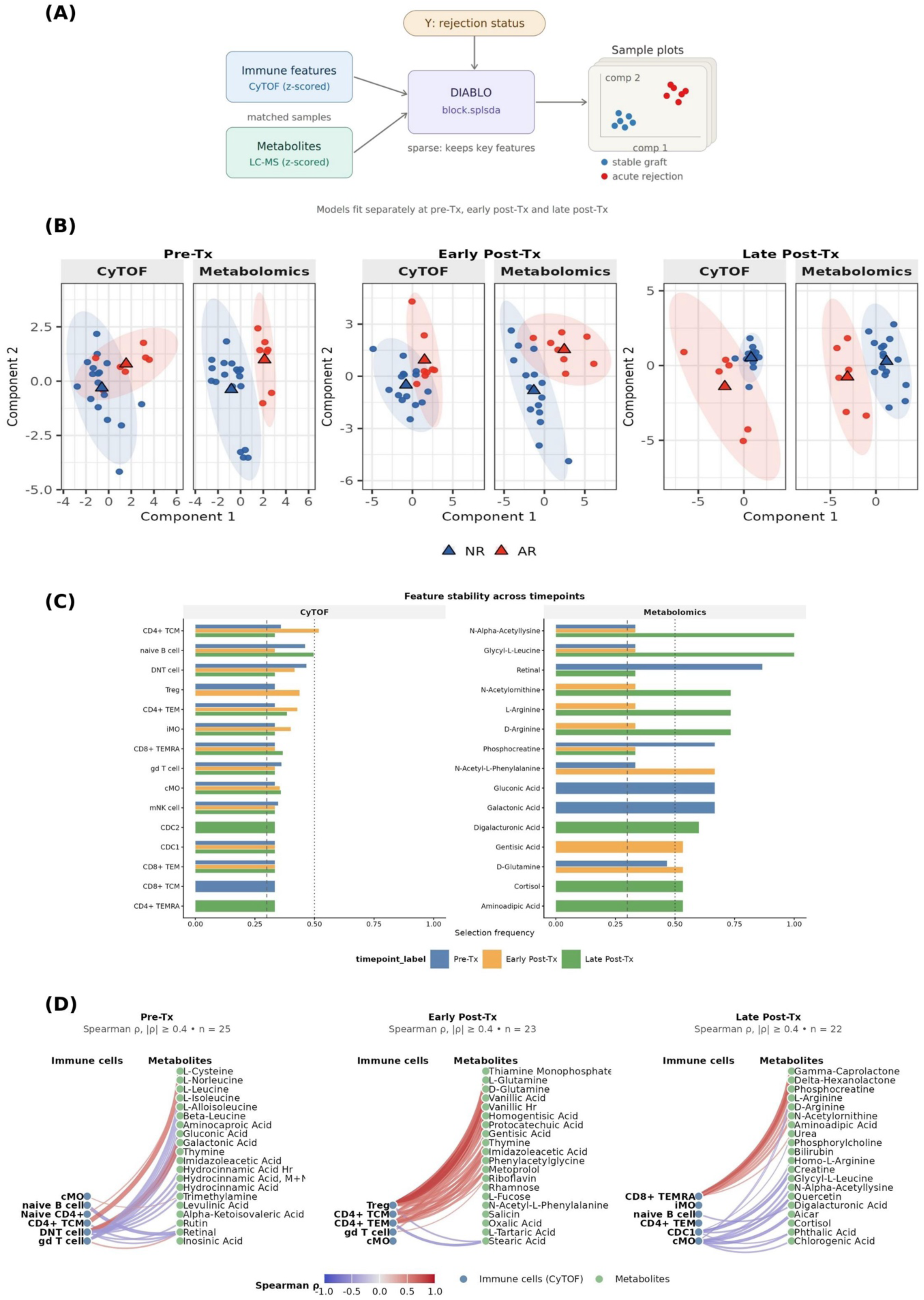
DIABLO multi-omics integration identifies timepoint-specific sample structure, feature stability, and immune–metabolite associations. **(A)** Schematic overview of DIABLO integration of matched CyTOF immune features and LC-MS metabolomics data using rejection status as the outcome. Models were fit separately at pre-, early post-, and late post-transplant timepoints. **(B)** DIABLO sample projections onto Components 1 and 2 for CyTOF and metabolomics blocks across timepoints. Each point represents one patient; triangles indicate group centroids; shaded ellipses indicate 95% confidence regions. NR, blue; AR, red. **(C)** Feature selection frequency across repeated 3-fold cross-validation of DIABLO model fitting. Dashed and dotted lines indicate 0.30 and 0.50 thresholds. **(D)** Spearman correlation networks between DIABLO-selected immune and metabolite features at each timepoint (|ρ| ≥ 0.4). Node color: blue, immune features; green, metabolites. Edge color: red, positive correlation; blue, negative correlation.

Cross-timepoint feature selection stability identified both recurring and temporally shifting contributors to DIABLO-defined discrimination (**Figure 9C**). Among CyTOF features, CD4+ TCM, DNT cells, naive B cells, Treg, and monocyte subsets were repeatedly selected across multiple timepoints, although their relative selection frequencies varied over the transplant course. Among metabolite features, N-alpha-acetyllysine, glycyl-L-leucine, retinal, N-acetylornithine, arginine-related metabolites, and phosphocreatine were among the most recurrently selected features, with additional metabolites showing timepoint-specific selection patterns. Several of these features map to a shared arginine-related axis, including L- and D-arginine, N-acetylornithine, creatine, and phosphocreatine, selected predominantly at early and late post-transplant. Ornithine and creatine are both downstream products of arginine metabolism, although these relationships were not tested directly in this dataset. Overall, these results suggest that immune and metabolic contributors to class separation varied across transplant stages.

To further examine cross-modal relationships among selected features, Spearman correlation networks were constructed at each timepoint using DIABLO-selected immune and metabolite features (|ρ| ≥ 0.4; **Figure 9D**). At pre-transplant, DNT cells showed the greatest number of edges with mixed positive and negative correlations across branched-chain amino acids, organic acids, and hydrocinnamic acid derivatives; CD4+ TCM contributed a predominantly positive correlation bundle across overlapping metabolites, while gd T cells showed broad negative correlations. Classical monocyte (cMO), naive B cell, and naive CD4+ T cells each connected to only a single metabolite. At early post-transplant, Tregs carried the densest positive correlation bundle spanning phenolic acids, glutamine species, thiamine monophosphate, and short-chain sugars; CD4+ TCM and CD4+ TEM contributed additional positive edges, and gd T cells showed sparse positive correlations. At late post-transplant, CD8+ TEMRA showed the broadest positive correlations with arginine-pathway metabolites, phosphocreatine, and organic acid lactones, while conventional dendritic cell type 1(cDC1) and cMO showed negative correlations with cortisol, phthalic acid, and chlorogenic acid; intermediate monocyte (iMO) and CD4+ TEM were absent from the network. The metabolites in this bundle included arginine, homoarginine, urea, creatine, and phosphocreatine, consistent with the recurrent selection of arginine-related features and with the established dependence of effector CD8+ T cell expansion on arginine and creatine availability(25), although these correlations remain exploratory. The cortisol association likely reflects corticosteroid exposure rather than endogenous biology and was not interpreted further. Collectively, these networks indicate a temporal shift in immune–metabolite co-variation patterns across the transplant course. Since these correlations were derived from features selected within a supervised integration framework, they should be considered exploratory associations.

## 4 Discussion

The ability to identify kidney transplant recipients at increased risk of rejection before clinical manifestation remains a major unmet need in transplantation. Using integrated transcriptomic, cytometric and metabolomic profiling across three independent and highly curated patient cohorts in which acute rejection samples were biopsy matched to confirm rejection phenotype, we identified a reproducible γδ T-cell effector signature associated with favorable allograft outcomes. Additionally, metabolomic features carried comparable discriminatory information, although no model was statistically superior in a cohort of this size. Importantly, γδ T-cell depletion associated with rejection was reproduced across three independent cohorts and orthogonal analytical platforms (RNA-sequencing, flow cytometry, and CyTOF), and in the cohort with pre-transplant sampling, this depletion was already detectable before transplantation and remained stable across the transplant course, suggesting that γδ T-cell biology reflects a component of recipients’ baseline immune state rather than a consequence of graft injury. Collectively, these findings identify a reproducible, mechanistically grounded γδ T-cell effector signature embedded within broader immune-metabolic programs associated with rejection susceptibility and raise γδ T-cell biology as a candidate direction for future pre-transplant risk assessment pending larger-scale validation.

γδ T-cells are a conserved, innate-like T-cell lineage that integrates stress sensing, immune surveillance and tissue repair. Their biology is well developed at the barrier sites, where resident γδ populations maintain integrity and respond rapidly to damage(10,45). Human γδ subsets are heterogeneous, with blood enriched for Vδ2-dominant effector populations, whereas tissue-compartments often contain more plastic Vδ1/ Vδ3 populations. In liver, γδ T-cells are clonally expanded and phenotypically distinct from matched blood populations, underscoring the organ context in shaping γδ states(13,46). In transplantation, the literature has suggested a dual role for γδ T-cells. In kidney allografts, γδ T-cells were found in rejected grafts, but not in non-rejecting kidneys, establishing an early link to rejection biology(17). Later work showed that CMV-responsive CD16+ γδ T-cells mediate antibody-dependent cellular cytotoxicity and are enriched in antibody-mediated kidney allograft lesions(18), whereas a rodent kidney model demonstrated that regulatory CD8+ γδ T-cells can prolong allograft survival via IL-10 linked suppression of alloimmunity(19). In contrast, in liver transplant, operational tolerance has been associated with Vδ1-skewed γδ signatures and even public clonotype selection, highlighting a tolerance-associated γδ trajectory that is much better established in liver than kidney(47). A transplant review stated that γδ T-cells can be pathogenic or protective depending on organ, subset, activation state, and microenvironment(48). What has remained unknown is whether a pre-transplant peripheral γδ state can forecast kidney transplant outcome before exposure to the allograft and immunosuppression. Our study addresses this gap: in a cohort sampled longitudinally, a specific γδ T-cell effector signature is already present before transplantation and is accompanied by metabolomic features associated with rejection; independently, γδ T-cell depletion associated with rejection is reproduced across transcriptomic, flow cytometry, and CyTOF platforms in three separate cohorts. In this sense, our kidney data begin to bridge the mechanistic gap between the liver literature, where γδ is tied to tolerance, and kidney literature, where γδ T cells have been interpreted as reactive or injury associated. Our data suggest that in kidney transplantation, a depleted mature γδ effector state may mark future rejection risk rather than simply reflect ongoing graft injury.

A notable strength of this study is the convergence of findings across independent biological layers. Transcriptomic analyses identified reduced γδ-associated TCR signatures in rejection and revealed enrichment of pathways related to antigen presentation, allograft rejection, and adaptive immune activation. These observations were reproduced in an independent US cohort and further refined through single-cell mass cytometry, which identified discrete γδ clusters depleted in future rejectors (**Figure 5**). The consistency of this pattern across various platforms substantially strengthens confidence that the observed signal reflects a genuine biological phenomenon. Reproducibility across orthogonal technologies is recognized as a critical benchmark for biomarker development in transplantation, where many promising signatures fail to generalize beyond discovery datasets. Importantly, the predictive γδ populations identified by CyTOF exhibited a mature effector phenotype characterized by elevated CD161, CD56, CD57, gdTCR, and T-bet expression and reduced expression of CD27 and CD45RA, markers associated with naïve differentiation states (**Supplementary Figure 5**). These observations suggest that the relevant biological signal may not be the numerical abundance per se but rather the presence of a specialized γδ effector compartment. CD161 has been linked to a conserved transcriptional program across T-cell lineages associated with innate-like cytokine responsiveness, mucosal tissue surveillance, and rapid TCR-independent effector functions(49,50). Similarly, T-bet, the master transcription factor of the IFN-γ-producing γδT1 lineage, identifies cytotoxic and tissue-surveillance-competent cells pre-programmed during thymic development(51).

The CD27-low/CD45RA-low phenotype is consistent with an effector-memory differentiation state, indicative of prior antigen exposure and the capacity for rapid recall responses, while remaining distinct from both naïve (CD45RA+CD27+) and terminally differentiated (CD45RA+CD27−) γδ T-cell populations(52,53). These γδ populations are enriched in non-rejectors before transplantation and remained detectable across time points, suggesting that they reflect a stable feature of recipient immune architecture. The selective depletion of this compartment in patients who later developed acute rejection therefore points to a pre-existing deficiency in γδ-mediated immune surveillance. This observation extends prior experimental evidence demonstrating that regulatory γδ T-cell subsets can suppress alloimmune responses and prolong kidney allograft survival through IL-10-dependent mechanism(19). Although the functional properties of the identified clusters remain to be determined, their phenotypic profile and longitudinal stability raise the possibility that loss of a mature, functionally poised γδ compartment may represent an early immunological vulnerability that predisposes to subsequent alloimmune injury.

Beyond immune-cell composition, our findings highlight the importance of immunometabolic regulation in transplantation. Mounting evidence indicates that immune cell function is tightly coupled to cellular metabolic programs and that metabolic rewiring influences differentiation, activation, and effector responses across multiple lymphocyte populations(54,55). In kidney transplantation, metabolomic studies have identified reproducible alterations in amino acid, carbohydrate, and lipid metabolism associated with acute rejection and graft dysfunction(56–61). Consistent with these observations, longitudinal metabolomic profiling revealed time-dependent metabolomic profiling associated with rejection status, with distinct pathway enrichment patterns observed across the pre, early-post, and late post-transplant periods (**Figure 6**, **Figure 7**). The most interpretable of these was enrichment of arginine and proline metabolism in acute rejection at early post-transplant, with arginine-related features including N-acetylornithine, creatine, and phosphocreatine independently selected in the integrative analysis and co-occurring with terminally differentiated CD8+ effector (TEMRA) cells at later timepoints. Given that arginine availability constrains T cell proliferation and effector function and that creatine supports CD8+ T cell expansion(23,25), this pattern is consistent with a metabolic context permissive to effector T cell responses, although the associations are correlative. Together with the other metabolic features identified, these observations suggest that γδ biology is embedded within a broader immunometabolic landscape rather than operating in isolation (**Figure 9**). The translational relevance of this study lies not in the identification of a single dominant biomarker, but in the convergence of independent immune and metabolic signals associated with rejection risk. While the models did not differ significantly in predictive performance **(Figure 8),** the immune and metabolic features were selected largely independently of one another, suggesting that these data layers capture complementary aspects of recipient biology. These discriminatory analyses (pre-transplant γδ alone, AUC = 0.69; metabolomics alone, AUC = 0.65; γδ combined with metabolomic features, AUC = 0.71) should be regarded as exploratory and hypothesis-generating given the sample size, rather than evidence of clinical-grade predictive performance. Given the absence of validated pre-transplant molecular tools for rejection risk in kidney transplantation, however, we view this as a candidate direction warranting future, adequately powered prospective validation. This finding aligns with an emerging view that transplant outcomes are shaped by coordinated immunological and metabolic programs rather than isolated cellular processes. As molecular monitoring strategies continue to evolve, integration of immune-cell phenotypes with metabolomic measurements may provide a complementary framework for risk stratification by capturing multiple dimensions of recipient state simultaneously. Although the predictive performance of the current models remains moderate, the biological concordance between the γδ T-cell and metabolomic signatures supports the existence of a broader immunometabolic landscape associated with susceptibility to rejection.

Several limitations warrant consideration. First, despite the use of independent cohorts and orthogonal validation approaches, overall sample size remains modest and may limit detection of more subtle biological effects. Second, of the three cohorts, only Cohort 3 included pre-transplant sampling; Cohorts 1 and 2 independently replicate γδ T-cell depletion in association with rejection using post-transplant, biopsy-matched samples, but do not themselves test whether this depletion precedes transplantation. Third, it remains unclear whether the identified γδ populations actively modulate alloimmunity or serve as a surrogate marker of broader immune competence. Fourth, γδ T-cell biology is influenced by CMV exposure, immunosuppressive regimen, age, inflammatory history and other variables that may contribute and should be examined systematically in the future. Fifth, metabolomic profiling was performed at all timepoints, but the interpretable metabolic signal emerged after transplantation, so the immune and metabolic layers are best viewed as complementary across the transplant course rather than as a single pre-transplant signature. Finally, prospective validation in a larger multicenter cohorts will be required before clinical implementation can be considered.

In summary, we identified a γδ T-cell depletion signature associated with kidney transplant outcome that is reproduced across independent cohorts and analytical platforms and, in a longitudinally sampled cohort, is detectable before transplantation and stable across the transplant course. Our findings suggest that γδ T-cell state is embedded within a broader immunometabolic landscape associated with susceptibility to rejection and illustrate the value of multi-omics approaches for resolving biologically relevant heterogeneity in transplant recipients.

## 5 Data Availability Statement

The datasets generated and/or analyzed during this study are not publicly available due to patient privacy considerations but are available from the corresponding author upon reasonable request.

## 6 Author Contributions

SMK and SS contributed equally and performed the analyses and wrote the manuscript. KK, SP, and TK contributed to data analysis and generation. OB and TKS provided samples and clinical data. MS, TH, JB, MHS, and MMS supervised the study and revised the manuscript. All authors approved the submitted version.

## 7 Funding

This project was supported in part by the National Institutes of Health/National Institute of Allergy and Infectious Diseases (NIH/NIAID), ancillary to U01AI063594 (CTOT21; PI: P.S. Heeger), with M.M. Sarwal as Principal Investigator of the ancillary CyTOF sub-study.

## 8 Conflict of Interest

MMS, SMK, SS, and MS are named inventors on a pending patent application related to the biomarkers described in this manuscript. The remaining authors declare that the research was conducted in the absence of any commercial or financial relationships that could be construed as a potential conflict of interest.

## Notes

### Competing Interest Statement

MMS, SMK, SS, and MS are named inventors on a pending provisional patent application related to the γδ T-cell and metabolomic biomarkers described in this manuscript. The remaining authors declare no competing interests.

## References

1. Tonelli M, Wiebe N, Knoll G, Bello A, Browne S, Jadhav D, et al. Systematic review: kidney transplantation compared with dialysis in clinically relevant outcomes. Am J Transplant Off J Am Soc Transplant Am Soc Transpl Surg. 2011 Oct;11(10):2093–109. doi:10.1111/j.1600-6143.2011.03686.x PubMed PMID: 21883901.

2. Lamb KE, Lodhi S, Meier-Kriesche HU. Long-term renal allograft survival in the United States: a critical reappraisal. Am J Transplant Off J Am Soc Transplant Am Soc Transpl Surg. 2011 Mar;11(3):450–62. doi:10.1111/j.1600-6143.2010.03283.x PubMed PMID: 20973913.

3. Ekberg H, Johansson ME. Challenges and considerations in diagnosing the kidney disease in deteriorating graft function. Transpl Int Off J Eur Soc Organ Transplant. 2012 Nov;25(11):1119–28. doi:10.1111/j.1432-2277.2012.01516.x PubMed PMID: 22738034; PubMed Central PMCID: PMC3487178.

4. Sigdel TK, Li L, Tran TQ, Khatri P, Naesens M, Sansanwal P, et al. Non-HLA antibodies to immunogenic epitopes predict the evolution of chronic renal allograft injury. J Am Soc Nephrol JASN. 2012 Apr;23(4):750–63. doi:10.1681/ASN.2011060596 PubMed PMID: 22302197.

5. Sarwal MM, Naesens M. Urine trumps the protocol biopsy for subclinical rejection surveillance. Kidney Int. 2023 Sep;104(3):432–9. doi:10.1016/j.kint.2023.06.023 PubMed PMID: 37599018.

6. Concepcion BP, Bestard O, Samaniego-Picota M, Prashar R, Ansari MJ, Comai G, et al. A Pre-Kidney Transplant Blood-Based Next-Generation Sequencing Assay to Predict Early Acute Rejection. Kidney360. 2026 May 14. doi:10.34067/KID.0000001208

7. Bestard O, Meneghini M, Crespo E, Bemelman F, Koch M, Volk HD, et al. Preformed T cell alloimmunity and HLA eplet mismatch to guide immunosuppression minimization with tacrolimus monotherapy in kidney transplantation: Results of the CELLIMIN trial. Am J Transplant. 2021 Aug;21(8):2833–45. doi:10.1111/ajt.16563

8. Hricik DE, Formica RN, Nickerson P, Rush D, Fairchild RL, Poggio ED, et al. Adverse Outcomes of Tacrolimus Withdrawal in Immune–Quiescent Kidney Transplant Recipients. J Am Soc Nephrol. 2015 Dec;26(12):3114–22. doi:10.1681/ASN.2014121234

9. Pineda S, Sigdel TK, Liberto JM, Vincenti F, Sirota M, Sarwal MM. Characterizing pre-transplant and post-transplant kidney rejection risk by B cell immune repertoire sequencing. Nat Commun. 2019 Apr 23;10(1):1906. doi:10.1038/s41467-019-09930-3 PubMed PMID: 31015506; PubMed Central PMCID: PMC6479061.

10. Bonneville M, O’Brien RL, Born WK. Gammadelta T cell effector functions: a blend of innate programming and acquired plasticity. Nat Rev Immunol. 2010 Jul;10(7):467–78. doi:10.1038/nri2781 PubMed PMID: 20539306.

11. Ribot JC, Lopes N, Silva-Santos B. γδ T cells in tissue physiology and surveillance. Nat Rev Immunol. 2021 Apr;21(4):221–32. doi:10.1038/s41577-020-00452-4 PubMed PMID: 33057185.

12. Chien Y hsiu, Meyer C, Bonneville M. γδ T cells: first line of defense and beyond. Annu Rev Immunol. 2014;32:121–55. doi:10.1146/annurev-immunol-032713-120216 PubMed PMID: 24387714.

13. Hunter S, Willcox CR, Davey MS, Kasatskaya SA, Jeffery HC, Chudakov DM, et al. Human liver infiltrating γδ T cells are composed of clonally expanded circulating and tissue-resident populations. J Hepatol. 2018 Sep;69(3):654–65. doi:10.1016/j.jhep.2018.05.007 PubMed PMID: 29758330; PubMed Central PMCID: PMC6089840.

14. Cerutti A, Cols M, Puga I. Marginal zone B cells: virtues of innate-like antibody-producing lymphocytes. Nat Rev Immunol. 2013 Feb;13(2):118–32. doi:10.1038/nri3383 PubMed PMID: 23348416; PubMed Central PMCID: PMC3652659.

15. Kaminski H, Marseres G, Yared N, Nokin MJ, Pitard V, Zouine A, et al. mTOR Inhibitors Prevent CMV Infection through the Restoration of Functional αβ and γδ T cells in Kidney Transplantation. J Am Soc Nephrol. 2022 Jan;33(1):121–37. doi:10.1681/ASN.2020121753

16. Marsères G, Gentil C, Tinevez C, Courant M, Cosentino A, Cornillot-Clément S, et al. Adoptive γδ T cell therapy controls cytomegalovirus infection in preclinical transplantation models. Nat Commun. 2026 Feb 17;17(1):2847. doi:10.1038/s41467-026-69538-2

17. Kirk AD, Ibrahim S, Dawson DV, Sanfilippo F, Finn OJ. Characterization of T cells expressing the gamma/delta antigen receptor in human renal allografts. Hum Immunol. 1993 Jan;36(1):11–9. doi:10.1016/0198-8859(93)90003-j PubMed PMID: 8458734.

18. Bachelet T, Couzi L, Pitard V, Sicard X, Rigothier C, Lepreux S, et al. Cytomegalovirus-responsive γδ T cells: novel effector cells in antibody-mediated kidney allograft microcirculation lesions. J Am Soc Nephrol JASN. 2014 Nov;25(11):2471–82. doi:10.1681/ASN.2013101052 PubMed PMID: 24744438; PubMed Central PMCID: PMC4214528.

19. Zhou J, Appleton SE, Stadnyk A, Lee TDG, Nashan BAP. CD8+ gammadelta T regulatory cells mediate kidney allograft prolongation after oral exposure to alloantigen. Transpl Int Off J Eur Soc Organ Transplant. 2008 Jul;21(7):679–87. doi:10.1111/j.1432-2277.2008.00669.x PubMed PMID: 18435686.

20. Wong G, Turner RM, Chapman JR, Howell M, Lim WH, Webster AC, et al. Time on dialysis and cancer risk after kidney transplantation. Transplantation. 2013 Jan 15;95(1):114–21. doi:10.1097/TP.0b013e31827743b4 PubMed PMID: 23222892.

21. Herrmann T, Fichtner AS, Karunakaran MM. An Update on the Molecular Basis of Phosphoantigen Recognition by Vγ9Vδ2 T Cells. Cells. 2020 Jun 9;9(6):1433. doi:10.3390/cells9061433 PubMed PMID: 32527033; PubMed Central PMCID: PMC7348870.

22. Lopes N, McIntyre C, Martin S, Raverdeau M, Sumaria N, Kohlgruber AC, et al. Distinct metabolic programs established in the thymus control effector functions of γδ T cell subsets in tumor microenvironments. Nat Immunol. 2021 Feb;22(2):179–92. doi:10.1038/s41590-020-00848-3

23. Rodriguez PC, Quiceno DG, Ochoa AC. L-arginine availability regulates T-lymphocyte cell-cycle progression. Blood. 2007 Feb 15;109(4):1568–73. doi:10.1182/blood-2006-06-031856 PubMed PMID: 17023580; PubMed Central PMCID: PMC1794048.

24. Munder M, Schneider H, Luckner C, Giese T, Langhans CD, Fuentes JM, et al. Suppression of T-cell functions by human granulocyte arginase. Blood. 2006 Sep 1;108(5):1627–34. doi:10.1182/blood-2006-11-010389 PubMed PMID: 16709924.

25. Di Biase S, Ma X, Wang X, Yu J, Wang YC, Smith DJ, et al. Creatine uptake regulates CD8 T cell antitumor immunity. J Exp Med. 2019 Dec 2;216(12):2869–82. doi:10.1084/jem.20182044 PubMed PMID: 31628186; PubMed Central PMCID: PMC6888972.

26. Xu T, Stewart KM, Wang X, Liu K, Xie M, Ryu JK, et al. Metabolic control of TH17 and induced Treg cell balance by an epigenetic mechanism. Nature. 2017 Aug 10;548(7666):228–33. doi:10.1038/nature23475 PubMed PMID: 28783731; PubMed Central PMCID: PMC6701955.

27. Barbetta A, Rocque B, Sarode D, Bartlett JA, Emamaullee J. Revisiting transplant immunology through the lens of single-cell technologies. Semin Immunopathol. 2023 Jan;45(1):91–109. doi:10.1007/s00281-022-00958-0 PubMed PMID: 35980400; PubMed Central PMCID: PMC9386203.

28. Kowli S, Minocherhomji S, Martinez OM, Busque S, Lebrec H, Maecker HT. Characterization of immune phenotypes in peripheral blood of adult renal transplant recipients using mass cytometry (CyTOF). ImmunoHorizons. 2025 Feb 18;9(4):vlae013. doi:10.1093/immhor/vlae013 PubMed PMID: 39965168; PubMed Central PMCID: PMC11841977.

29. Kuypers DRJ, Kamphorst JJ, Loor H de, O’Day EM. Perspective: metabolomics has the potential to change the landscape of kidney transplantation diagnostics. Biomark Med. 2024;18(17– 18):787–94. doi:10.1080/17520363.2024.2394383 PubMed PMID: 39234983; PubMed Central PMCID: PMC11457662.

30. Pineda S, Sur S, Sigdel T, Nguyen M, Crespo E, Torija A, et al. Peripheral Blood RNA Sequencing Unravels a Differential Signature of Coding and Noncoding Genes by Types of Kidney Allograft Rejection. Kidney Int Rep. 2020 Oct;5(10):1706–21. doi:10.1016/j.ekir.2020.07.023 PubMed PMID: 33102963; PubMed Central PMCID: PMC7569686.

31. Vivian J, Rao AA, Nothaft FA, Ketchum C, Armstrong J, Novak A, et al. Toil enables reproducible, open source, big biomedical data analyses. Nat Biotechnol. 2017 Apr 11;35(4):314–6. doi:10.1038/nbt.3772 PubMed PMID: 28398314; PubMed Central PMCID: PMC5546205.

32. Bolotin DA, Poslavsky S, Mitrophanov I, Shugay M, Mamedov IZ, Putintseva EV, et al. MiXCR: software for comprehensive adaptive immunity profiling. Nat Methods. 2015 May;12(5):380–1. doi:10.1038/nmeth.3364 PubMed PMID: 25924071.

33. R Development Core Team. R: A language and environment for statistical computing. [Internet]. Vienna, Austria: R Foundation for Statistical Computing; 2008. Available from: https://www.r-project.org/

34. Dobin A, Gingeras TR. Mapping RNA-seq Reads with STAR. Curr Protoc Bioinforma. 2015 Sep 3;51:11.14.1–11.14.19. doi:10.1002/0471250953.bi1114s51 PubMed PMID: 26334920; PubMed Central PMCID: PMC4631051.

35. Liu JX, Gao YL, Xu Y, Zheng CH, You J. Differential expression analysis on RNA-Seq count data based on penalized matrix decomposition. IEEE Trans Nanobioscience. 2014 Mar;13(1):12–8. doi:10.1109/TNB.2013.2296978 PubMed PMID: 24594510.

36. Love MI, Huber W, Anders S. Moderated estimation of fold change and dispersion for RNA-seq data with DESeq2. Genome Biol. 2014;15(12):550. doi:10.1186/s13059-014-0550-8 PubMed PMID: 25516281; PubMed Central PMCID: PMC4302049.

37. Zou H, Hastie T. Regularization and Variable Selection Via the Elastic Net. J R Stat Soc Ser B Stat Methodol. 2005 Apr 1;67(2):301–20. doi:10.1111/j.1467-9868.2005.00503.x

38. Chen EY, Tan CM, Kou Y, Duan Q, Wang Z, Meirelles GV, et al. Enrichr: interactive and collaborative HTML5 gene list enrichment analysis tool. BMC Bioinformatics. 2013 Apr 15;14:128. doi:10.1186/1471-2105-14-128 PubMed PMID: 23586463; PubMed Central PMCID: PMC3637064.

39. Fabregat A, Jupe S, Matthews L, Sidiropoulos K, Gillespie M, Garapati P, et al. The Reactome Pathway Knowledgebase. Nucleic Acids Res. 2018 Jan 4;46(D1):D649–55. doi:10.1093/nar/gkx1132 PubMed PMID: 29145629; PubMed Central PMCID: PMC5753187.

40. Croft D, Mundo AF, Haw R, Milacic M, Weiser J, Wu G, et al. The Reactome pathway knowledgebase. Nucleic Acids Res. 2014 Jan;42(Database issue):D472–477. doi:10.1093/nar/gkt1102 PubMed PMID: 24243840; PubMed Central PMCID: PMC3965010.

41. Genuer R, Poggi JM, Tuleau-Malot C. VSURF: An R Package for Variable Selection Using Random Forests. R J. 2015;7(2):19. doi:10.32614/RJ-2015-018

42. Kanehisa M, Goto S. KEGG: kyoto encyclopedia of genes and genomes. Nucleic Acids Res. 2000 Jan 1;28(1):27–30. doi:10.1093/nar/28.1.27 PubMed PMID: 10592173; PubMed Central PMCID: PMC102409.

43. Korotkevich G, Sukhov V, Budin N, Shpak B, Artyomov MN, Sergushichev A. Fast gene set enrichment analysis [Internet]. Bioinformatics; 2016 [cited 2026 Jul 20]. Available from: http://biorxiv.org/lookup/doi/10.1101/060012 doi:10.1101/060012

44. Singh A, Shannon CP, Gautier B, Rohart F, Vacher M, Tebbutt SJ, et al. DIABLO: an integrative approach for identifying key molecular drivers from multi-omics assays. Bioinformatics. 2019 Sep 1;35(17):3055–62. doi:10.1093/bioinformatics/bty1054 PubMed PMID: 30657866; PubMed Central PMCID: PMC6735831.

45. Nielsen MM, Witherden DA, Havran WL. γδ T cells in homeostasis and host defence of epithelial barrier tissues. Nat Rev Immunol. 2017 Dec;17(12):733–45. doi:10.1038/nri.2017.101 PubMed PMID: 28920588; PubMed Central PMCID: PMC5771804.

46. Vantourout P, Hayday A. Six-of-the-best: unique contributions of γδ T cells to immunology. Nat Rev Immunol. 2013 Feb;13(2):88–100. doi:10.1038/nri3384 PubMed PMID: 23348415; PubMed Central PMCID: PMC3951794.

47. Zhao X, Li Y, Ohe H, Nafady-Hego H, Uemoto S, Bishop GA, et al. Intragraft Vδ1 γδ T cells with a unique T-cell receptor are closely associated with pediatric semiallogeneic liver transplant tolerance. Transplantation. 2013 Jan 15;95(1):192–202. doi:10.1097/TP.0b013e3182782f9f PubMed PMID: 23222896.

48. Sullivan LC, Shaw EM, Stankovic S, Snell GI, Brooks AG, Westall GP. The complex existence of γδ T cells following transplantation: the good, the bad and the simply confusing. Clin Transl Immunol. 2019;8(9):e1078. doi:10.1002/cti2.1078 PubMed PMID: 31548887; PubMed Central PMCID: PMC6748302.

49. Fergusson JR, Smith KE, Fleming VM, Rajoriya N, Newell EW, Simmons R, et al. CD161 defines a transcriptional and functional phenotype across distinct human T cell lineages. Cell Rep. 2014 Nov 6;9(3):1075–88. doi:10.1016/j.celrep.2014.09.045 PubMed PMID: 25437561; PubMed Central PMCID: PMC4250839.

50. Karunathilaka A, Halstrom S, Price P, Holt M, Lutzky VP, Doolan DL, et al. CD161 expression defines new human γδ T cell subsets. Immun Ageing A. 2022 Feb 22;19(1):11. doi:10.1186/s12979-022-00269-w PubMed PMID: 35193613; PubMed Central PMCID: PMC8862246.

51. Fiala GJ, Gomes AQ, Silva-Santos B. From thymus to periphery: Molecular basis of effector γδ-T cell differentiation. Immunol Rev. 2020 Nov;298(1):47–60. doi:10.1111/imr.12918 PubMed PMID: 33191519; PubMed Central PMCID: PMC7756812.

52. Dieli F, Poccia F, Lipp M, Sireci G, Caccamo N, Di Sano C, et al. Differentiation of effector/memory Vdelta2 T cells and migratory routes in lymph nodes or inflammatory sites. J Exp Med. 2003 Aug 4;198(3):391–7. doi:10.1084/jem.20030235 PubMed PMID: 12900516; PubMed Central PMCID: PMC2194087.

53. Odaira K, Kimura SN, Fujieda N, Kobayashi Y, Kambara K, Takahashi T, et al. CD27(-)CD45(+) γδ T cells can be divided into two populations, CD27(-)CD45(int) and CD27(-)CD45(hi) with little proliferation potential. Biochem Biophys Res Commun. 2016 Sep 23;478(3):1298–303. doi:10.1016/j.bbrc.2016.08.115 PubMed PMID: 27553282.

54. Egawa T, Bhattacharya D. Regulation of metabolic supply and demand during B cell activation and subsequent differentiation. Curr Opin Immunol. 2019 Apr;57:8–14. doi:10.1016/j.coi.2018.10.003

55. Hu C, Xuan Y, Zhang X, Liu Y, Yang S, Yang K. Immune cell metabolism and metabolic reprogramming. Mol Biol Rep. 2022 Oct;49(10):9783–95. doi:10.1007/s11033-022-07474-2

56. Mao YY, Bai JQ, Chen JH, Shou ZF, He Q, Wu JY, et al. A pilot study of GC/MS-based serum metabolic profiling of acute rejection in renal transplantation. Transpl Immunol. 2008 Apr;19(1):74–80. doi:10.1016/j.trim.2008.01.006 PubMed PMID: 18346641.

57. O’Neill LAJ, Kishton RJ, Rathmell J. A guide to immunometabolism for immunologists. Nat Rev Immunol. 2016 Sep;16(9):553–65. doi:10.1038/nri.2016.70 PubMed PMID: 27396447; PubMed Central PMCID: PMC5001910.

58. Soyer T, Özyüksel G, Türer ÖB, Çakmakkaya K, Yavuz S, Yalçın B, et al. Bilateral Pulmonary Langerhans’s Cell Histiocytosis is Surgical Challenge in Children: A Case Report. Eur J Pediatr Surg Rep. 2019 Jan;7(1):e8–11. doi:10.1055/s-0039-1688771 PubMed PMID: 31131184; PubMed Central PMCID: PMC6533102.

59. Gen S, Wright AC. Policy Capacity Is Necessary but Not Sufficient Comment on “Health Reform Requires Policy Capacity.” Int J Health Policy Manag. 2015 Aug 2;4(12):837–9. doi:10.15171/ijhpm.2015.145 PubMed PMID: 26673469; PubMed Central PMCID: PMC4663088.

60. Vander Heiden MG, DeBerardinis RJ. Understanding the Intersections between Metabolism and Cancer Biology. Cell. 2017 Feb 9;168(4):657–69. doi:10.1016/j.cell.2016.12.039 PubMed PMID: 28187287; PubMed Central PMCID: PMC5329766.

61. Zengeler KE, Lukens JR. Innate immunity at the crossroads of healthy brain maturation and neurodevelopmental disorders. Nat Rev Immunol. 2021 Jul;21(7):454–68. doi:10.1038/s41577-020-00487-7 PubMed PMID: 33479477; PubMed Central PMCID: PMC9213174.

