## Supplementary Materials for "A Reproducible Pre-Transplant γδ T-cell Effector Signature Reveals Immune-Metabolic Programs Associated with Kidney Allograft Rejection"

**Supplemental Table 1. Genes associated with the αβ/γδ TCR ratio identified by elastic net regression.**

| No. | Gene | Direction in rejection |
|---|---|---|
| 1 | ADRB2 | Downregulated |
| 2 | APOBEC3G | Downregulated |
| 3 | C2 | Downregulated |
| 4 | C9orf43 | Downregulated |
| 5 | CCL4 | Downregulated |
| 6 | CCR5 | Downregulated |
| 7 | CD8B | Downregulated |
| 8 | CRIM1 | Downregulated |
| 9 | EXO1 | Downregulated |
| 10 | FASLG | Downregulated |
| 11 | GZMA | Downregulated |
| 12 | GZMB | Downregulated |
| 13 | GZMH | Downregulated |
| 14 | IPP | Downregulated |
| 15 | KLHL4 | Downregulated |
| 16 | KLRG1 | Downregulated |
| 17 | PSME2 | Downregulated |
| 18 | R3HCC1L | Downregulated |
| 19 | RP11-26J3.4 | Downregulated |
| 20 | SOCS4 | Downregulated |
| 21 | SUPV3L1 | Downregulated |
| 22 | TBX21 | Downregulated |
| 23 | VPS41 | Downregulated |
| 24 | ABHD4 | Upregulated |
| 25 | AL049794.1 | Upregulated |
| 26 | ATXN7L2 | Upregulated |
| 27 | C15orf41 | Upregulated |
| 28 | CDH26 | Upregulated |
| 29 | CRIP2 | Upregulated |
| 30 | DLX4 | Upregulated |
| 31 | FAM196B | Upregulated |
| 32 | GARNL3 | Upregulated |
| 33 | GCHFR | Upregulated |
| 34 | GEMIN8 | Upregulated |
| 35 | GGT5 | Upregulated |
| 36 | GYLTL1B | Upregulated |
| 37 | HLF | Upregulated |
| 38 | HOXB3 | Upregulated |

| No. | Gene | Direction in rejection |
|---|---|---|
| 39 | ICA1L | Upregulated |
| 40 | ITIH4 | Upregulated |
| 41 | KDM5B | Upregulated |
| 42 | LYVE1 | Upregulated |
| 43 | MAFF | Upregulated |
| 44 | MAP7D1 | Upregulated |
| 45 | MCOLN3 | Upregulated |
| 46 | METTL12 | Upregulated |
| 47 | MFSD6L | Upregulated |
| 48 | MGRN1 | Upregulated |
| 49 | MTSS1L | Upregulated |
| 50 | MYBPH | Upregulated |
| 51 | NPC2 | Upregulated |
| 52 | NPTN | Upregulated |
| 53 | NTRK3 | Upregulated |
| 54 | PER3 | Upregulated |
| 55 | PGC | Upregulated |
| 56 | PPARGC1A | Upregulated |
| 57 | PPP5D1 | Upregulated |
| 58 | PRKCZ | Upregulated |
| 59 | RBM20 | Upregulated |
| 60 | RFX2 | Upregulated |
| 61 | RNF41 | Upregulated |
| 62 | RSRP1 | Upregulated |
| 63 | SAMD14 | Upregulated |
| 64 | SCRG1 | Upregulated |
| 65 | SGTA | Upregulated |
| 66 | SLC22A14 | Upregulated |
| 67 | SRGAP3 | Upregulated |
| 68 | TMEM200A | Upregulated |
| 69 | TPBG | Upregulated |
| 70 | TRIM47 | Upregulated |
| 71 | UBAC2 | Upregulated |
| 72 | UPB1 | Upregulated |
| 73 | WDR83 | Upregulated |
| 74 | ZC3H12B | Upregulated |
| 75 | ZFHX2 | Upregulated |
| 76 | ZGLP1 | Upregulated |
| 77 | ZNF76 | Upregulated |

| No. | Gene | Direction in rejection |
|---|---|---|
| 78 | ZNF774 | Upregulated |
| 79 | ZNHIT1 | Upregulated |

**Supplementary Table 2. FDR-adjusted p-values for FlowSOM clusters ranked by PLS-DA VIP score**

Results for the top 20 FlowSOM clusters selected by PLS-DA variable importance in projection (VIP) score, comparing cluster frequencies between acute rejection (AR) and non-rejection (NR) patients. P-values adjusted using the Benjamini-Hochberg (BH) procedure.

| Cluster | Cell Subset | FDR-adjusted p-value (BH) |
|---|---|---|
| 18 | mNK cell | 0.1560 |
| 5 | mNK cell | 0.1560 |
| 17 | mNK cell | 0.2850 |
| 172 | iNK cell | 0.2768 |
| 78 | iNK cell | 0.1560 |
| 91 | iNK cell | 0.1403 |
| 64 | iNK cell | 0.1403 |
| 52 | γδ T cell | 0.0196 |
| 55 | γδ T cell | 0.1403 |
| 66 | γδ T cell | 0.0196 |
| 67 | γδ T cell | 0.0196 |
| 26 | CD8+ TEMRA | 0.2024 |
| 215 | cMO | 0.1560 |
| 211 | cMO | 0.1560 |
| 152 | cMO | 0.1612 |
| 154 | cMO | 0.1560 |
| 167 | cMO | 0.2700 |
| 141 | DNT cell | 0.1560 |
| 163 | naive B cell | 0.1560 |
| 177 | naive B cell | 0.1560 |

**Supplementary Table 3. Cross-validated classification performance of timepoint-specific DIABLO models.**

Balanced error rates for the CyTOF and metabolomics blocks on components 1 and 2, estimated by repeated M-fold cross-validation (3 folds, 5 repeats) using the centroids distance. Values below 0.5 indicate performance exceeding chance for two-class classification.

| Timepoint | CyTOF BER (c1, c2) | Metabolomics BER (c1, c2) |
|---|---|---|
| Pre-Transplant | 0.416, 0.400 | 0.352, 0.336 |
| Early post-transplant | 0.435, 0.504 | 0.400, 0.435 |
| Late post-transplant | 0.409, 0.345 | 0.255, 0.255 |

Averaged Over # of Samples

| | STA | CMR | AMR |
| --- | --- | --- | --- |
| IGH Reads | 366 | 176 | 285 |
| IGK Reads | 436 | 204 | 439 |
| IGL Reads | 197 | 108 | 221 |
| TRA Reads | 4,647 | 3,765 | 6,311 |
| TRB Reads | 1,127 | 573 | 611 |
| TRD Reads | 160 | 63 | 61 |
| TRG Reads | 301 | 170 | 163 |
| **Total Reads** | **7,234** | **5,059** | **8,090** |
| IGH Clones | 6 | 4 | 4 |
| IGK Clones | 85 | 39 | 77 |
| IGL Clones | 23 | 13 | 22 |
| TRA Clones | 143 | 63 | 75 |
| TRB Clones | 164 | 81 | 79 |
| TRD Clones | 8 | 2 | 3 |
| TRG Clones | 68 | 34 | 26 |
| **Total Clones** | **497** | **237** | **286** |

Define a clone as: same V, J, CDR3 length and 90% identity of the CDR3 region

**Supplemental Figure 1. Immune repertoire composition by transplant outcome in Cohort 1 (Bulk RNA-seq, n=36).**

TCR and BCR immune repertoire reads and clonotypes were extracted from bulk RNA-seq data using MiXCR, aligning RNA-seq fragments to VDJ genomic regions. A clone was defined as cells sharing identical V gene, J gene, CDR3 length, and ≥90% CDR3 amino acid sequence identity. Values represent means averaged across samples within each outcome group: AMR (n=12), TCMR (n=13), and STA (n=11). Reads and clonotypes are reported separately for immunoglobulin heavy and light chains (IGH, IGK, IGL) and TCR chains (TRA, TRB, TRD, TRG). TRD and TRG encode the γδ TCR chains; reduced TRD and TRG reads and clonotypes in TCMR and AMR relative to STA are consistent with γδ T cell depletion in rejecting allografts. Total reads and total clonotypes represent the sum across all seven receptor loci per group.

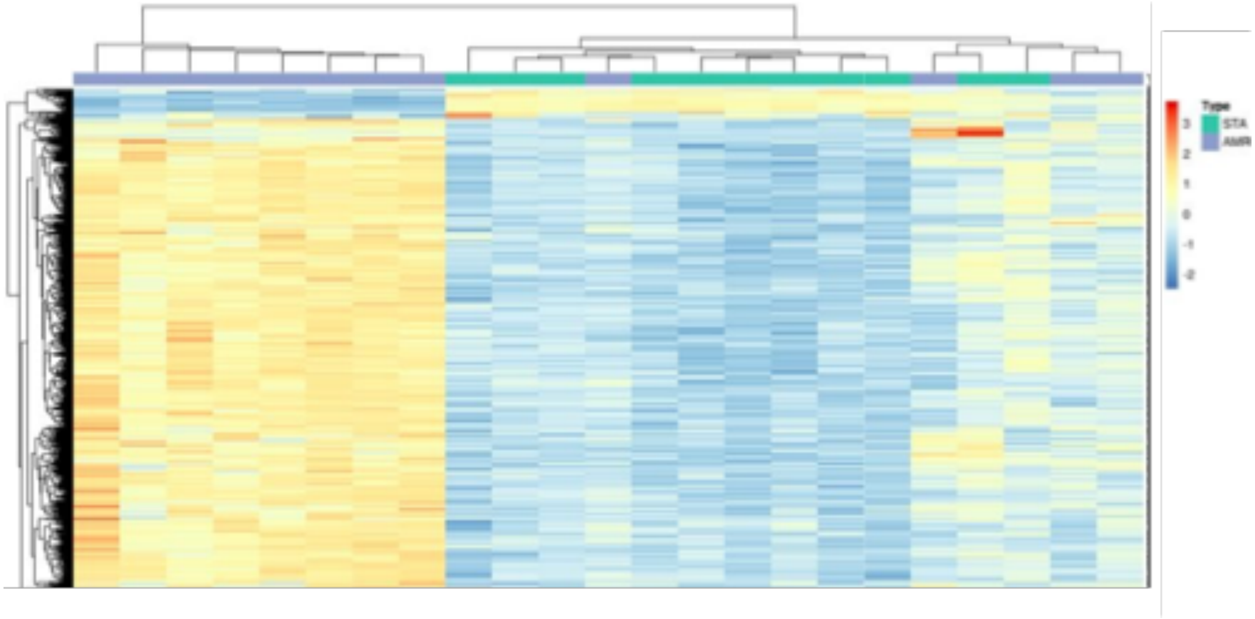


**Supplemental Figure 2. Heatmap showcasing differential gene expression between STA and AMR samples.**

Horizontal axis represents each individual sample (with color indicating transplant outcome); vertical axis lists the 1716 significantly differentially expressed genes. Of these 1716 genes, 1506 are upregulated and 210 are downregulated in the AMR samples.

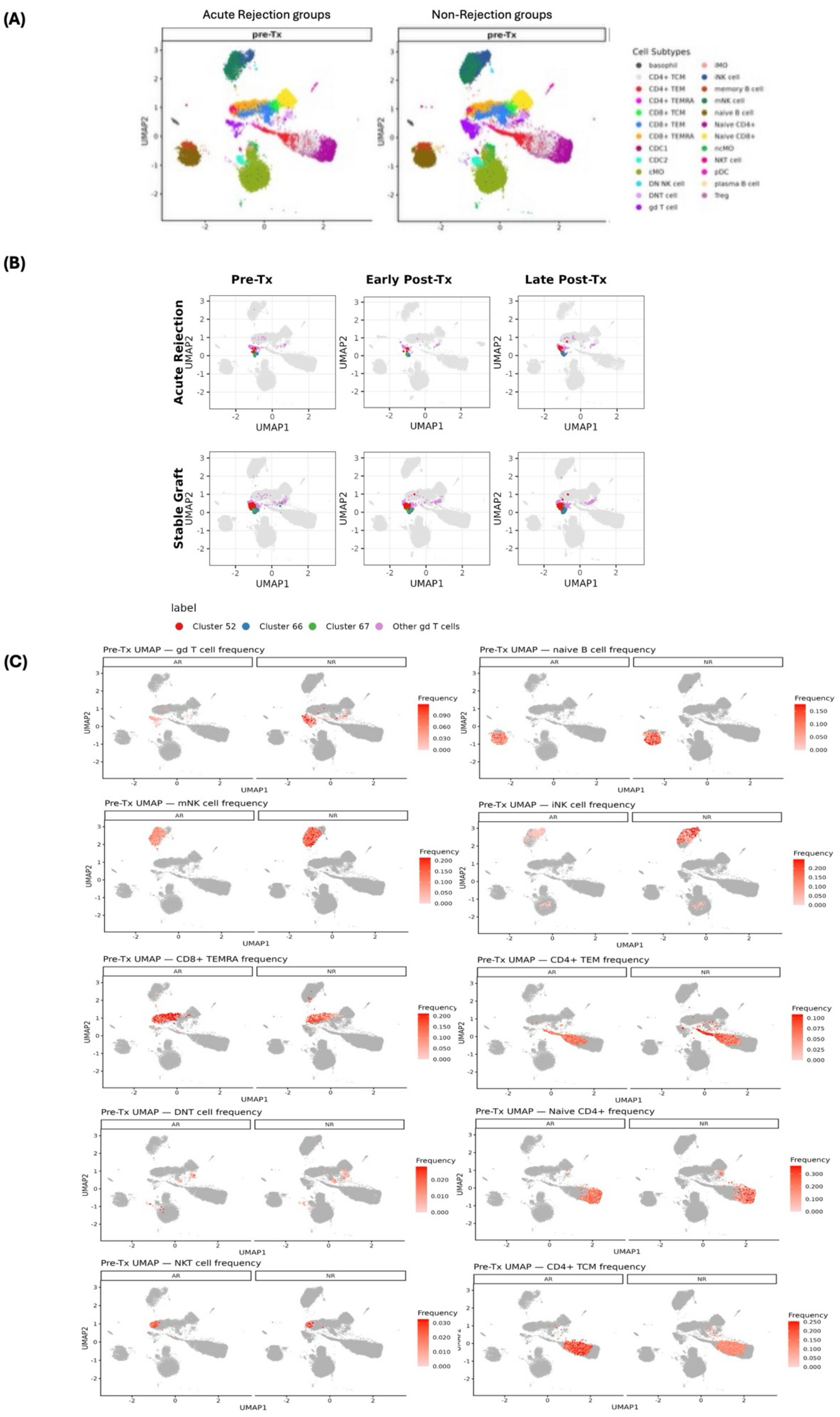

(A)
Acute Rejection groups
Non-Rejection groups
pre-Tx
Cell Subtypes
basophil
CD4+ TCM
CD4+ TEM
CD4+ TEMRA
CD8+ TCM
CD8+ TEM
CD8+ TEMRA
CDC1
CDC2
cMO
DN NK cell
DNT cell
gd T cell
iMO
iNK cell
memory B cell
mNK cell
naive B cell
Naive CD4+
Naive CD8+
ncMO
NKT cell
pDC
plasma B cell
Treg
UMAP2
(B)
Pre-Tx
Early Post-Tx
Late Post-Tx
Acute Rejection
Stable Graft
UMAP1
UMAP2
label
Cluster 52
Cluster 66
Cluster 67
Other gd T cells
(C)
Pre-Tx UMAP — gd T cell frequency
Pre-Tx UMAP — naive B cell frequency
Pre-Tx UMAP — mNK cell frequency
Pre-Tx UMAP — iNK cell frequency
Pre-Tx UMAP — CD8+ TEMRA frequency
Pre-Tx UMAP — CD4+ TEM frequency
Pre-Tx UMAP — DNT cell frequency
Pre-Tx UMAP — Naive CD4+ frequency
Pre-Tx UMAP — NKT cell frequency
Pre-Tx UMAP — CD4+ TCM frequency
AR
NR
Frequency
UMAP1
UMAP2

**Supplementary Figure 3. Pre-transplant immune landscape reveals baseline differences between acute rejection and non-rejection patients.**

**(A)** Pre-transplant peripheral immune cells from acute rejection and non-rejection patients share a broadly similar global architecture when visualized by UMAP, suggesting that outcome-associated differences are subtle rather than wholesale compositional shifts. **(B)** γδ T cell clusters 52, 66, and 67 mapped across all timepoints (pre-transplant, early post-transplant, late post-transplant), shown separately for acute rejection and stable graft patients. All three clusters occupy a tight, overlapping UMAP region, reflecting phenotypically similar γδ T cell states. **(C)** Pre-transplant UMAP feature plots for the top immune subsets selected at pre-transplant by PLS-DA illustrate their spatial localization across acute rejection and non-rejection patients. These plots are shown as qualitative visualization of the PLS-DA–prioritized features, while quantitative interpretation is based on the corresponding patient-level frequency analyses. AR, acute rejection; NR, stable graft.

### a) Pre-transplant immune cell subset abundance

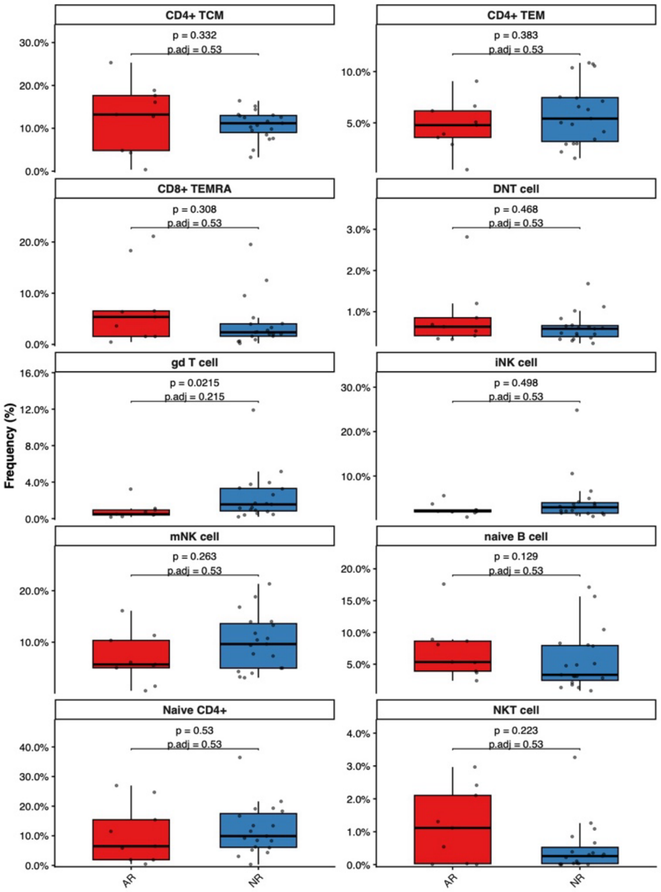


### b) Early post-transplant immune cell subset abundance

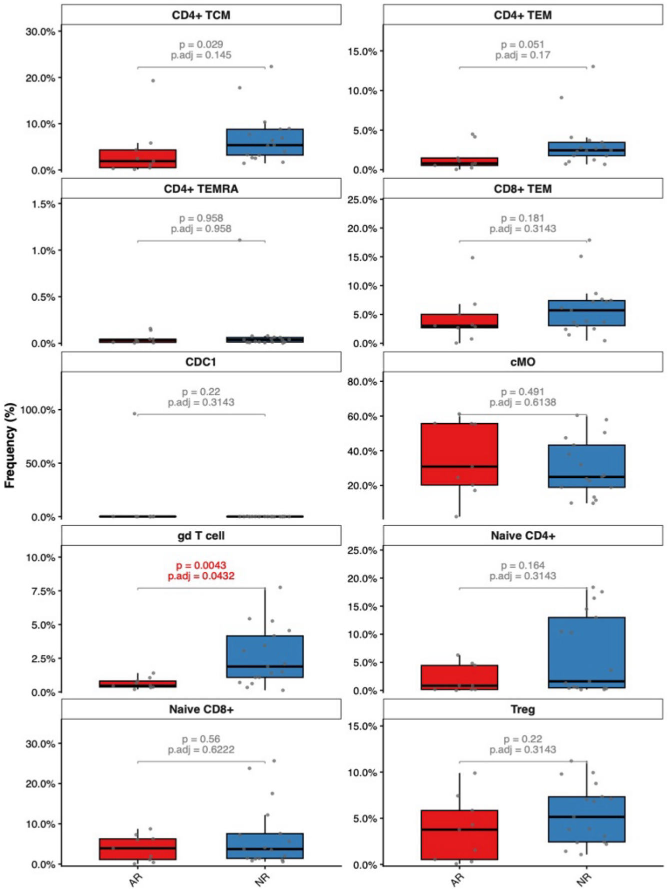

**Supplementary Figure 4. Pre- and early post-transplant abundance of PLS-DA–prioritized immune cell subsets**

Immune cell subsets were prioritized based on PLS-DA variable importance in projection (VIP) scores, calculated as the combined VIP across the first two PLS-DA components; the top 10 subsets by VIP score were carried forward for testing. **(A)** Box plots comparing pre-transplant frequencies of the top immune cell subsets ranked by PLS-DA VIP score between acute rejection (AR) and non-rejection (NR) patients. Each point represents an individual patient. Raw p-values and FDR-adjusted p-values (BH correction) are displayed for each subset. **(B)** Box plots comparing early post-transplant frequencies of the top immune cell subsets ranked by PLS-DA VIP score between AR and NR patients. Each point represents one patient. Raw and BH-adjusted p-values are shown; γδ T cells were significantly reduced in AR patients after FDR correction.

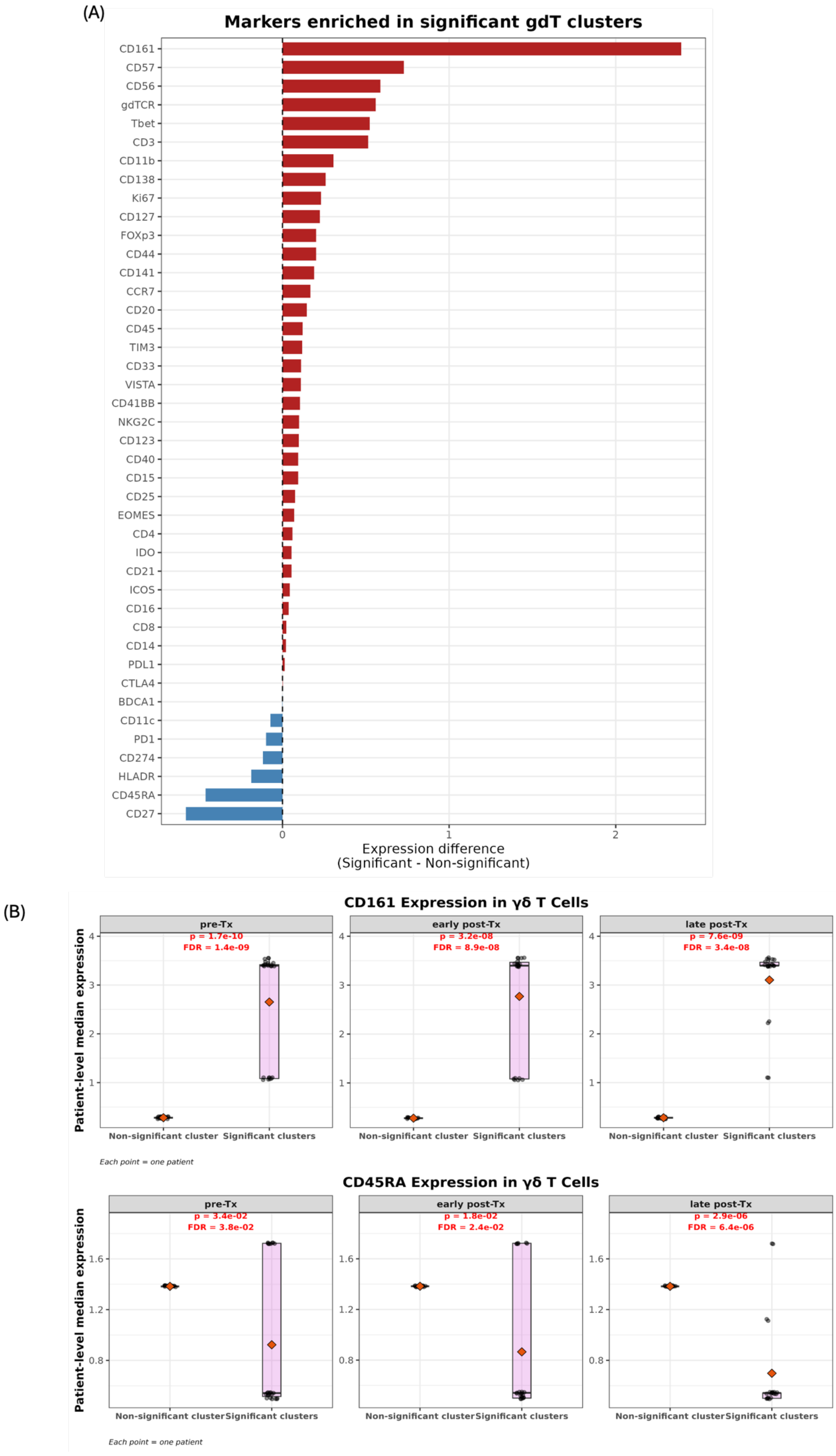
(A)
Markers enriched in significant gdT clusters
CD161
CD57
CD56
gdTCR
Tbet
CD3
CD11b
CD138
Ki67
CD127
FOXp3
CD44
CD141
CCR7
CD20
CD45
TIM3
CD33
VISTA
CD41BB
NKG2C
CD123
CD40
CD15
CD25
EOMES
CD4
IDO
CD21
ICOS
CD16
CD8
CD14
PDL1
CTLA4
BDCA1
CD11c
PD1
CD274
HLADR
CD45RA
CD27
0
1
2
Expression difference
(Significant - Non-significant)
(B)
CD161 Expression in γδ T Cells
pre-Tx
p = 1.7e-10
FDR = 1.4e-09
early post-Tx
p = 3.2e-08
FDR = 8.9e-08
late post-Tx
p = 7.6e-09
FDR = 3.4e-08
Patient-level median expression
Non-significant cluster
Significant clusters
Each point = one patient
CD45RA Expression in γδ T Cells
pre-Tx
p = 3.4e-02
FDR = 3.8e-02
early post-Tx
p = 1.8e-02
FDR = 2.4e-02
late post-Tx
p = 2.9e-06
FDR = 6.4e-06
Patient-level median expression
Non-significant cluster
Significant clusters
Each point = one patient

**Supplementary Figure 5. Marker enrichment profile of significant γδ T cell clusters.**

**(A)** Differential marker expression between the three significant γδ T cell clusters (52, 66, 67) and cluster 55. Significant clusters are broadly enriched for activated and cytotoxic markers — most prominently CD161, CD57, CD56, gdTCR, and T-bet — while showing reduced expression of naïve-associated markers CD27 and CD45RA, suggesting a mature, terminally differentiated effector phenotype. **(B)** Median expression of the two most divergent markers, CD161 and CD45RA, in γδ T cells at pre-transplant, early post-transplant, and late post-transplant timepoints, compared between the non-significant cluster (55) and the three significant clusters (52, 66, 67). Both markers differ significantly between the cluster groups at every timepoint after correction (BH-adjusted FDR $< 0.05$), with CD161 enriched and CD45RA reduced in the significant clusters, consistent with the mature effector phenotype described above.
